# Spatially-local inhibition and synaptic plasticity together enable dynamic, context-dependent integration of parallel sensory pathways

**DOI:** 10.1101/2025.10.18.683230

**Authors:** Qiang Chen, Fred Rieke

## Abstract

Retinal ganglion cells have traditionally been grouped into cells that are sensitive to luminance but not spatial structure and cells with responses that are enhanced by spatial structure. Neither category captures responses of mouse Off Transient alpha cells, which are largest for spatially homogeneous inputs and are suppressed by spatial structure. We identified two circuit mechanisms that together can explain this unusual spatial selectivity. First, inhibition to these cells is tuned to finer spatial structure than excitation, causing the balance of excitation and inhibition to depend on spatial scale. Second, the excitatory synapses onto these cells undergo strong synaptic depression and the modulation of that depression by presynaptic inhibition amplifies responses to the transition from spatially structured to homogeneous inputs. A spatiotemporal computational model incorporating these circuit features quantitatively recapitulates the observed dynamics. These findings reveal how localized inhibition and short-term plasticity jointly create the distinctive spatial selectivity of Off Transient cells.

## Introduction

Interactions between excitation and inhibition (E/I) are a cornerstone of neural computation. Excitatory and inhibitory inputs are often dynamically balanced and stimulus-dependent modulation of this balance shapes sensitivity of sensory neurons to specific stimulus features (Monier et al., 2003; Murphy and Rieke, 2006; Cardin et al., 2007; Okun and Lampl, 2008; Cafaro and Rieke, 2010; Isaacson and Scanziani, 2011). While many studies focus on changes in E/I balance, less is known about how they interact with other circuit mechanisms to shape circuit computation. Here we investigate how E/I balance and short-term synaptic plasticity together create an unusual selectivity for spatially homogeneous visual inputs that differentiates one retinal ganglion cell type from its counterparts.

Inhibitory circuits shape stimulus selectivity in several ways. Inhibition in many sensory circuits is more broadly tuned (e.g. in visual space or across odors) than excitation (Wu et al., 2008; Poo and Isaacson, 2009). The resulting lateral inhibition sharpens stimulus selectivity by amplifying the differences in responses of neurons that are strongly and weakly activated by a given stimulus. Inhibitory circuits often play a similar sharpening role in the time domain, where temporally-delayed inhibition can sharpen temporal tuning by creating a narrow window of opportunity for spiking (Pouille and Scanziani, 2001; Wehr and Zador, 2003; Murphy and Rieke, 2006). Excitation and inhibition are often finely balanced and hence subtle differences in the amplitude or timing of inhibition relative to excitation can have large effects on neural responses. E/I interactions occur in the context of other circuit mechanisms such as short-term synaptic plasticity (Anwar et al., 2017; Deng et al., 2024). These circuit mechanisms can alter the E/I balance and impact computation. In the hippocampus, for example, excitatory and inhibitory signals exhibit opposite forms of plasticity and the resulting dynamic changes in E/I balance can enhance signals characteristic of place fields (Klyachko and Stevens, 2006; Rotman et al., 2011; Anwar et al., 2017). In the retina, short term plasticity at inhibitory synapses can lead to selective suppression of responses to repeated stimuli and enhancement of those to novel stimuli (Hosoya et al., 2005).

Our goal here was to exploit knowledge of how signals are routed through retinal circuits to study the general issue of how E/I balance works in concert with other circuit mechanisms to control circuit outputs. Retinal ganglion cells (RGCs) typically respond more strongly to spatially structured inputs than to unstructured inputs. Off-transient alpha RGCs (OffT αRGCs) are an exception - responding more strongly to spatially homogeneous inputs. We find that this preference for homogenous inputs emerges from the interaction of two distinct circuit features. First, inhibitory signals are tuned to finer spatial structure than excitatory signals, causing inhibition to preferentially cancel responses to fine spatial structure. This is the opposite of well-studied lateral inhibition (Hartline et al., 1956; Olsen and Wilson, 2008). Second, the excitatory synapses onto Off-transient cells exhibit strong synaptic depression. Presynaptic inhibition recruited by spatially structured inputs relieves this depression, causing strong responses to subsequent spatially-homogeneous inputs.

## Results

We start by identifying and characterizing two circuit mechanisms that contribute to the selectivity of OffT αRGCs for spatially homogeneous inputs. First, we show that the inhibitory circuits that control OffT responses are sensitive to smaller spatial scales than the excitatory inputs - causing inhibition and excitation to preferentially cancel for fine spatial scale inputs. This originates from the divergence of common inputs to two parallel pathways with distinct spatial pooling properties. Second, we show that presynaptic inhibition relieves synaptic depression at excitatory synapses and by doing so enhances responses to spatially homogeneous inputs. Finally, we construct a quantitative model incorporating these two mechanisms and show that it can replicate key features of the OffT αRGC responses.

Our studies build on prior knowledge of the three parallel pathways that convey rod signals through the rodent retina under these conditions (Völgyi et al., 2004; Manookin et al., 2008; Grimes et al., 2014, 2018; Ke et al., 2014; Jin et al., 2022) (Bloomfield and Dacheux, 2001; Völgyi et al., 2004; Grimes et al., 2018; Jin et al., 2022) (Figure 1A; Supplementary Figure 1): (1) the primary rod bipolar → A2 amacrine pathway, (2) the secondary rod-cone gap junction pathway, and (3) the tertiary direct rod-to-off cone bipolar cell pathway. At the mesopic light levels we study here (~100 R*/rod/s), all three pathways are active and converge onto OffT αRGC, conditions under which their preference for homogeneous inputs is strong. We leverage this understanding of rod pathway organization to investigate how interactions between these parallel circuits shape the distinctive spatial selectivity of OffT cells.

**Figure 1.**
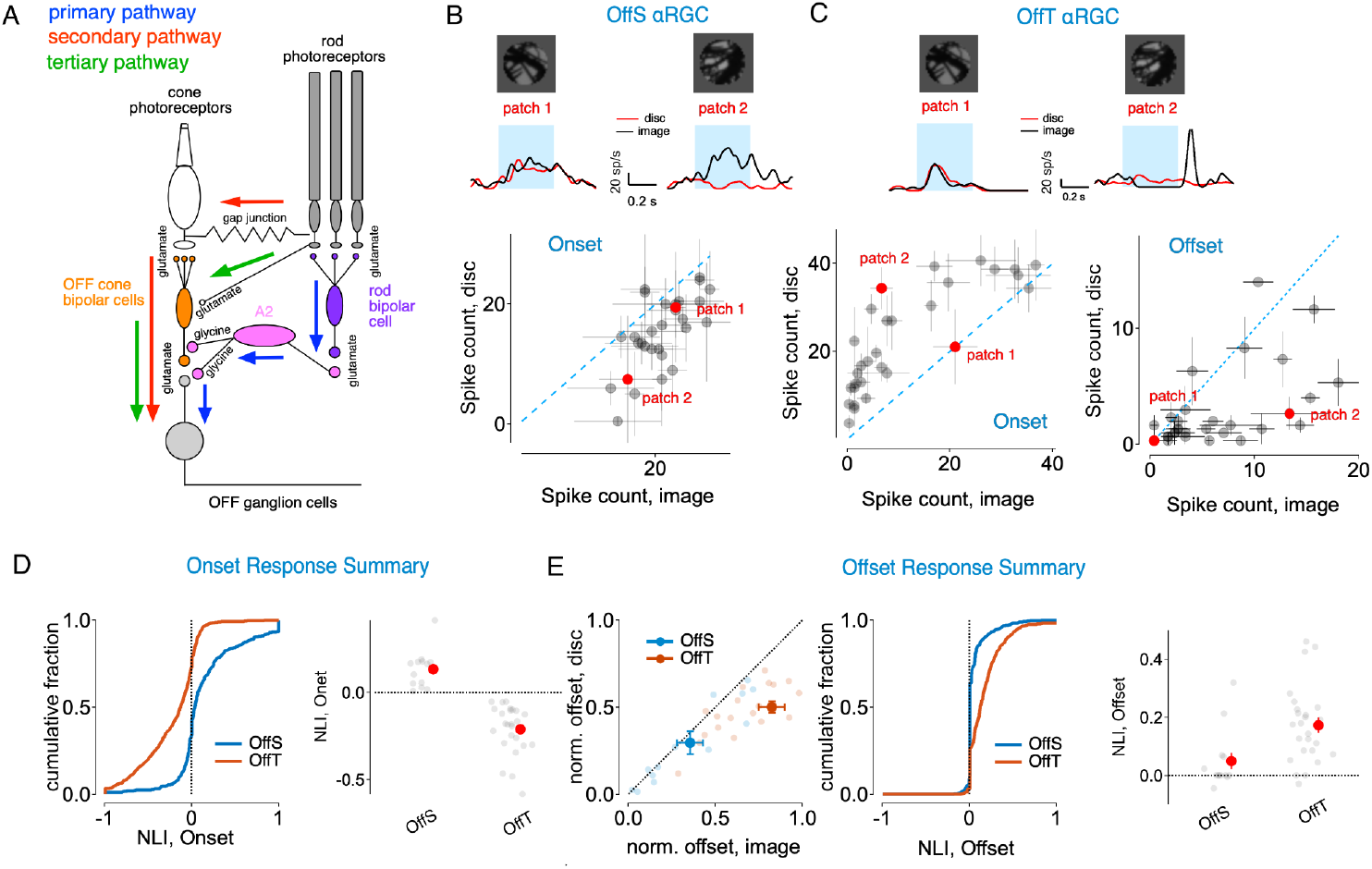
Nonlinear Spatial Integration Across αRGCs. (A) Schematics for parallel rod signaling pathways. The primary pathway (blue arrows) routes rod signals through rod bipolar cells and AII amacrine cells to ganglion cells. The secondary pathway (pink arrows) utilizes rod-cone gap junctions, allowing rod signals to flow through cone bipolar cells to ganglion cells. The tertiary pathway (green arrows) directly connects rods to OFF cone bipolar cells, activating OFF ganglion cells. (B) Spatial integration in Off-sustained (OffS) alpha retinal ganglion cells. Top: Example spike responses to two image patches and their corresponding equivalent discs. Bottom: Scatter plot for onset response, indicating preference for spatially structured stimuli. Error bars represent SEM. (C) Spatial integration in Off-transient (OffT) alpha retinal ganglion cells. Top: Example spike responses to two image patches (black traces) and their corresponding linear-equivalent discs (red traces) during stimulus presentation (blue shading). Bottom: Scatter plots comparing spike counts for image patches versus equivalent discs for onset (left) and offset (right) responses. Error bars represent SEM. (D) Onset response summary. Left: Cumulative distribution of onset nonlinearity indices across all image-patch/disc pairs for OffS (blue) and OffT (orange) cells. Note that OffT cells predominantly show negative NLIs. Right: Summary of mean onset NLIs for individual cells (gray circles) and population means+S.E.M. (red circles) for OffS and OffT types. (E) Offset response summary. Left: offset responses normalized to the mean of onset responses, comparing responses to linear-equivalent discs versus natural image patches for all alpha RGC subtypes. This normalization reveals that OffT cells (orange) exhibit substantially stronger offset responses relative to other cell types. Middle: Cumulative distribution of offset nonlinearity indices for OffS and OffT cells. Right: summary of mean offset NLIs for individual cells (gray circles) and population means (red circles) for OffS and OffT types. Note: when offset response is <3 spikes for both disc and image, the offset NLI is set to 0.

**Supplementary Figure 1.**
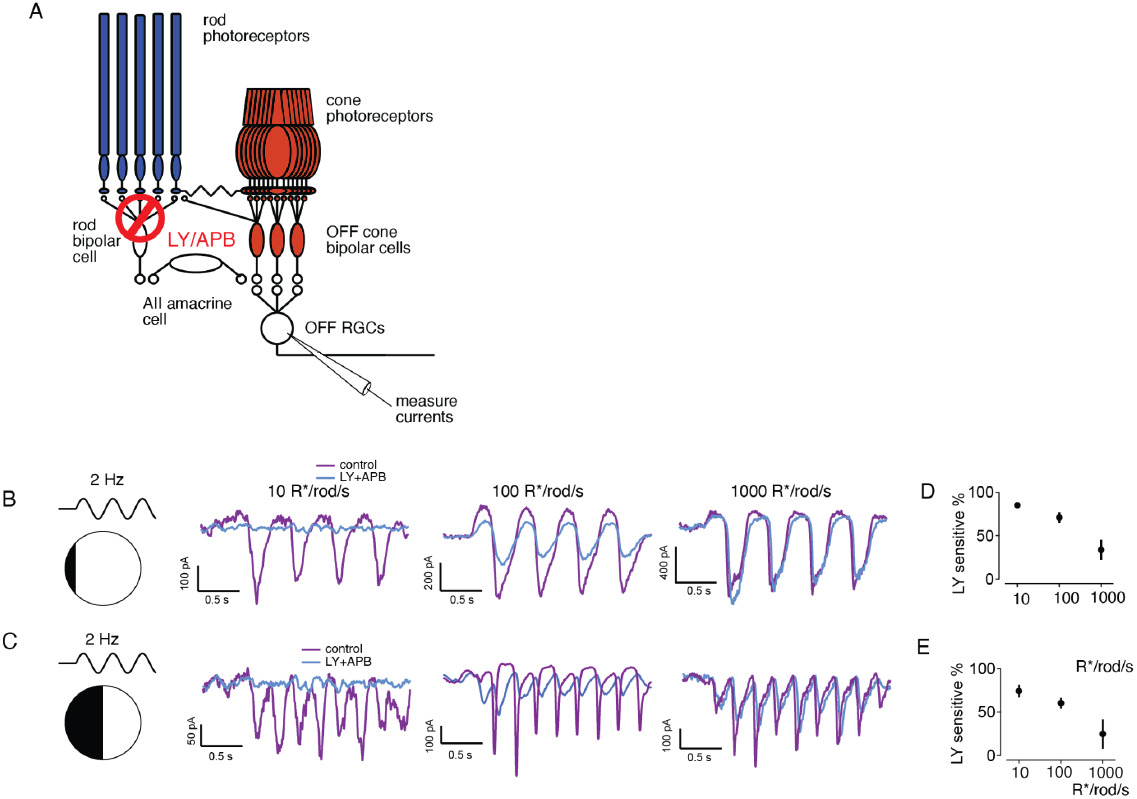
Routing of Rod-derived signals under different light levels. (A) Schematic of the experimental setup showing pharmacological isolation of rod pathways using LY/APB to block rod bipolar cell signaling in the primary rod pathway. (B) Example current traces (control, purple; LY+APB treatment, blue) from OffT αRGC in response to a 2 Hz full-field modulating spots at 10 R*/rod/s, 100 R*/rod/s, and 1000 R*/rod/s. (C) Similar current traces showing OffT αRGC responses to a contrast-reversing grating stimulus. (D-E) Quantification of the LY/APB-sensitive component of the OffT αRGC responses at different light intensities (R*/rod/s).

### Off-Transient αRGCs Exhibit Robust Homogeneity Preference at Stimulus Onset and Offset

We assessed how RGCs integrated spatial inputs within their receptive field (RF) center by comparing responses to flashed natural image patches with responses to spatially is homogeneous ‘linear-equivalent’ discs (Turner and Rieke, 2016; Turner et al., 2018)(Figure 1 B-C). All stimuli were confined to the RF center estimated for each cell at the start of recording (see Methods). The intensity of each linear-equivalent disc was calculated as a weighted linear sum of the pixel intensities in the corresponding image patch, with weights taken from a Gaussian profile matching the cell’s RF center(Turner and Rieke, 2016; Turner et al., 2018). If a RGC integrated spatial inputs linearly, responses to the natural image patch and its corresponding linear-equivalent disc should be near identical. Substantial disparities in these responses thus indicated nonlinear spatial integration.

When we compared responses to natural images versus homogeneous discs, we found distinct nonlinear spatial integration characteristics among Off αRGC subtypes (Figure 1B-E). At stimulus onset, Off-Sustained (OffS) αRGCs typically exhibited enhanced responses to spatially structured natural image patches compared to linear-equivalent discs (Figure 1B). Off-Transient (OffT) αRGCs, however, consistently showed stronger responses to linear-equivalent discs at stimulus onset, indicating a preference for spatial homogeneity (Figure 1C, left). OffT, but not OffS, αRGCs also exhibited robust responses to the transition from a spatially structured to a uniform image (i.e. at stimulus offset; Figure 1C, right). This highlights the sensitivity of these cells to spatially uniform inputs.

We quantified the deviation from linear spatial integration for each image patch using a nonlinearity index (NLI), defined as

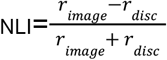

Here *r*_*image*_ and *r*_*disc*_ are the average spike counts in response to the original natural image patch and the corresponding linear-equivalent disc. The NLI quantifies different aspects of spatial selectivity at stimulus onset versus offset. At stimulus onset, positive NLI values indicate a preference for spatial structure, whereas negative NLIs indicate a preference for spatial homogeneity. At offset, the NLI measures responses to the transition from structured images back to uniform background: positive values indicate homogeneity preference—cells respond more strongly when returning from spatial structure to homogeneity. NLIs for OffT cells were significantly negative at onset and significantly positive at offset relative to OffS αRGCs (and other αRGCs subtypes; Figure 1D-E).

Together, the findings summarized in Figure 1 demonstrate that OffT αRGCs exhibit a robust homogeneity preference. These distinctive responses of OffT αRGCs to the onset and offset of spatially-structured stimuli differentiate them from other αRGC subtypes.

### Synaptic Mechanisms of OffT αRGC Homogeneity Preference

Having established the distinctive homogeneity preference of OffT αRGCs, we next investigated the underlying synaptic mechanisms mediating these responses using naturalistic stimuli and controlled artificial stimuli.

We recorded excitatory and inhibitory synaptic inputs to OffT αRGCs in response to natural image patches and their linear-equivalent discs (Figure 2). At stimulus onset, excitatory inputs to OffT αRGCs demonstrated a clear preference for homogeneous discs, evident from the preponderance of data points that fall above the unity line and the negative onset NLIs (Figure 2A-C). Spatial structure often elicited a reduction in tonic or baseline excitatory current compared to homogeneous discs (e.g. patch 1 in Figure 2A)—suggestive of presynaptic inhibition that was preferentially recruited by spatially-structured stimuli. Conversely, inhibitory synaptic inputs displayed stronger responses to structured natural images compared to linear-equivalent discs, indicated by data points falling below the unity line and positive onset NLIs (Figure 2E-G). These findings suggest that spatial structure in natural images robustly activates inhibitory pathways, likely involving both direct inhibition to OffT αRGCs and presynaptic inhibition to the bipolar cells that provide excitatory input to OffT cells.

**Figure 2.**
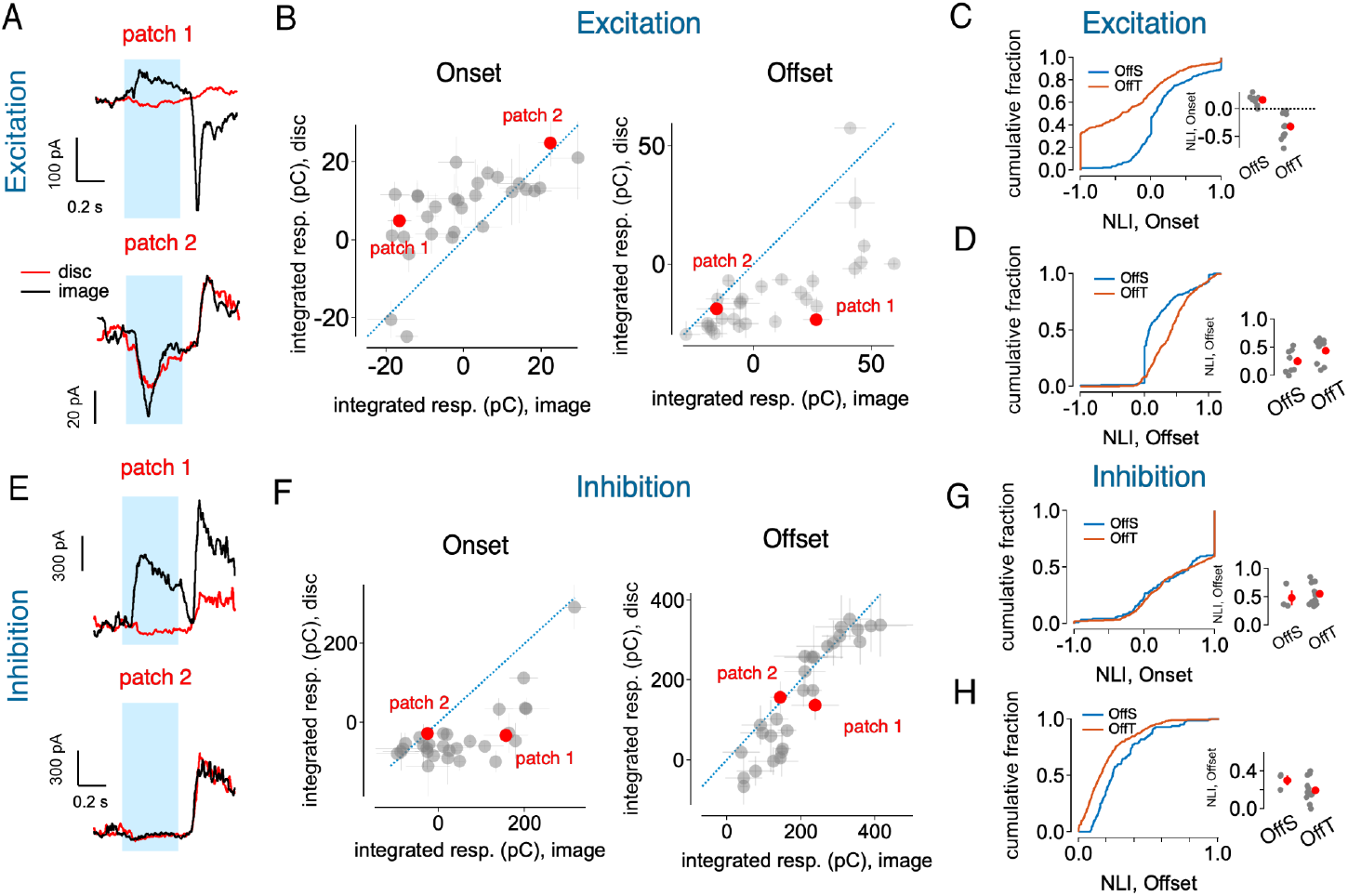
Synaptic mechanisms underlying OffT αRGC responses to natural images. (A) Example excitatory current traces recorded from an OffT αRGC in response to two natural image patches (black) and their corresponding linear-equivalent discs (red). The blue shaded region indicates the stimulus presentation period. (B) Scatter plots comparing excitatory input charge transfer (pC) in response to natural image patches versus linear-equivalent discs. Left: At stimulus onset, excitatory inputs show stronger responses to linear-equivalent discs than to natural image patches (points above the unity line). Right: At stimulus offset, excitatory inputs show robust responses to the transition from natural images to a uniform background. (C) Cumulative distributions of onset nonlinearity indices (NLIs) for excitatory inputs across different αRGC types. Left: Distribution curves for OffS (blue) and OffT (orange) cells. Right: Summary of mean excitatory onset NLIs for individual cells (gray circles) and population means +S.E.M (red circles). (D) Cumulative distributions of offset nonlinearity indices for excitatory inputs in OffS (blue) and OffT (orange) cells. Left: Distribution curves. Right: Mean offset NLIs for individual cells (gray circles) and population means +S.E.M (red circles). (E) Example inhibitory current traces recorded from an OffT αRGC in response to two natural image patches (black) and their corresponding linear-equivalent discs (red). (F) Scatter plots comparing inhibitory input (left: onset; right: offset) charge transfer (pC) in response to natural image patches versus linear-equivalent discs. (G) Cumulative distributions of onset nonlinearity indices for inhibitory inputs across different αRGC types. Left: Distribution curves for OffS (blue) and OffT (orange) cells. Right: Summary of mean inhibitory onset NLIs for individual cells (gray circles) and population means +S.E.M (red circles). (H) same as G but for offset responses.

The return from a structured image to spatial homogeneity at stimulus offset often substantially increased excitatory synaptic inputs to OffT αRGCs (Figure 2B,D), consistent with the positive offset NLI previously observed in the spike response. Patches that elicited a robust increase in excitatory synaptic input at stimulus offset also showed a clear suppression of excitatory input at stimulus onset (e.g. patch 1 in Figure 2).

To more precisely isolate the impact of spatial structure, we conducted additional experiments using flashed gratings with a range of bar sizes (Figure 3). The onset of a flashed grating strongly suppressed OffT αRGC spike responses through a pronounced suppression of baseline excitatory input and an increase in inhibitory input (Figure 3A-C). At grating offset, OffT αRGCs exhibited robust spike responses characterized by a marked increase in excitatory input (Figure 3B). These temporal response profiles differed from those of OffS αRGCs, for which excitatory and inhibitory synaptic inputs increased at both grating onset and offset (Figure 3D-F). This distinction emphasizes an unexpected difference in the synaptic mechanisms controlling responses of these two Off αRGC types.

**Figure 3:**
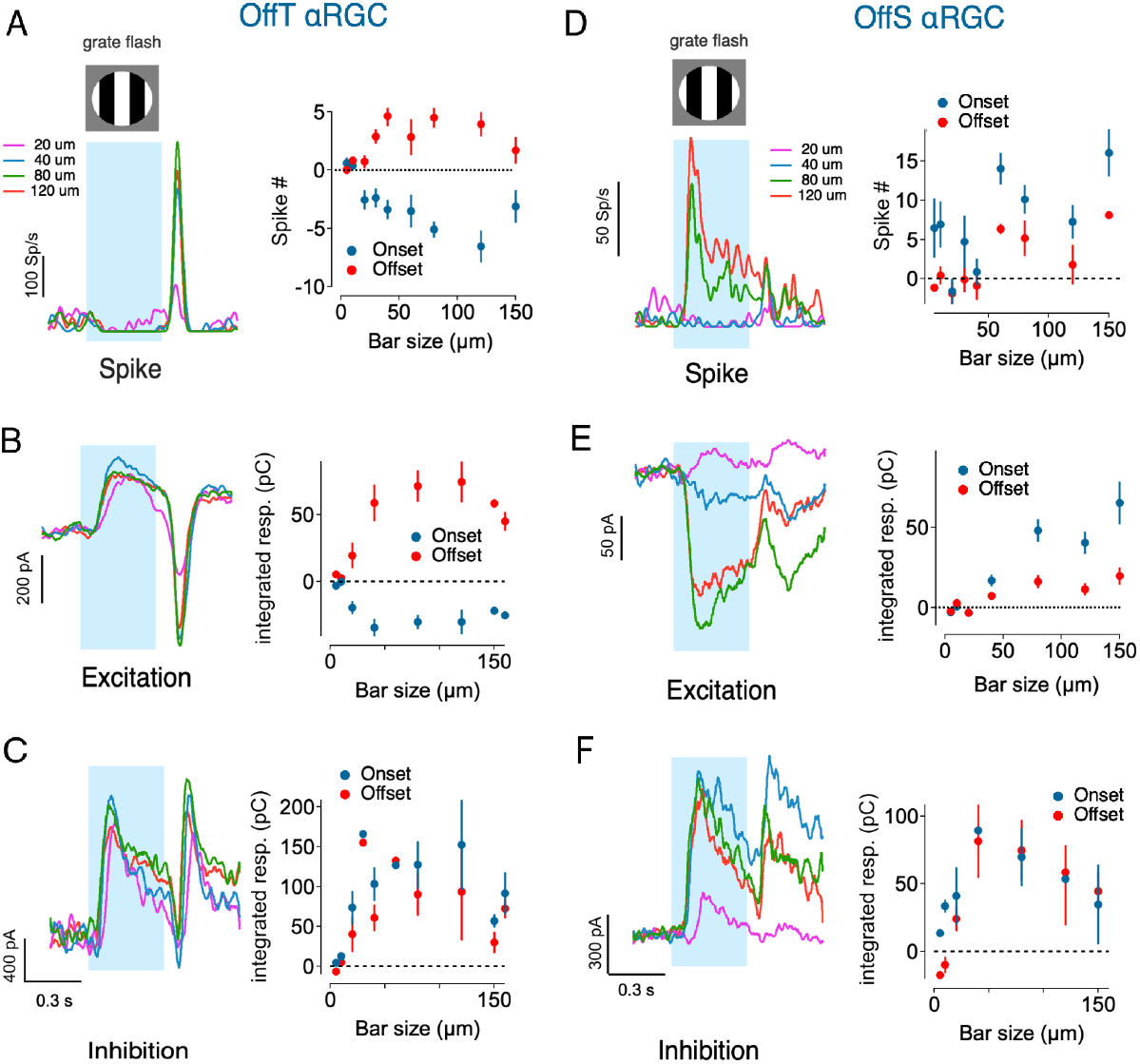
Comparison of synaptic responses to flashed gratings between OffT and OffS αRGCs. (A) Response properties of OffT alpha retinal ganglion cells to flashed gratings. Left panel shows example spike traces for four different bar sizes with blue shading indicating the flash duration. Right panel shows quantified spike responses as a function of bar size for both onset (blue circles) and offset (red circles) of the flash. Note that negative spike responses at onset indicate suppression of spike relative to baseline firing. (B) Excitatory synaptic input recordings from OffT αRGCs in response to flashed gratings. Left panel shows example excitatory current traces for the four bar sizes. Right panel shows the integrated excitatory response as a function of bar size at both onset (blue) and offset (red). Excitatory responses exhibit suppression during grating presentation and enhancement at stimulus offset. (C) Inhibitory synaptic input recordings from OffT αRGCs in response to flashed gratings. Left panel shows example inhibitory current traces for the four bar sizes. Right panel shows the integrated inhibitory response as a function of bar size at both onset (blue) and offset (red). (D-F) same as A-C, but for OffS αRGCs.

Together, these experiments point to potential circuit mechanisms that could generate the OffT αRGCs’ unusual homogeneity preference. First, spatially-structured stimuli appear to recruit inhibition both directly on OffT αRGCs and presynaptically on their bipolar inputs, as evidenced by the suppression of baseline excitation. Second, the pronounced rebound at the offset of spatial structure hints at a gain control mechanism. The experiments below probe each of these mechanisms in more detail.

### Spatially Localized Inhibition Creates Scale-Dependent E/I Balance in OffT αRGCs

We next studied interactions between excitation and inhibition. We start by examining their spatial-temporal properties in more detail. At the mesopic light levels we studied, excitation and inhibition are routed through parallel pathways, with inhibitory synaptic inputs to Off αRGCs derived predominantly through the primary pathway and excitatory inputs derived from a mixture of primary and secondary pathways (Figure 1A) (Ke et al., 2014; Grimes et al., 2018). This parallel routing suggested that the properties of the nonlinear receptive field subunits that control responses to spatial structure might differ for excitatory and inhibitory synaptic inputs to Off αRGCs.

We characterized the subunit properties of excitatory and inhibitory inputs using whole-cell voltage clamp recordings while employing a classic test for nonlinear subunits using contrast-reversing gratings (Hochstein and Shapley, 1976). In the case of linear spatial integration, responses to the bright and dark bars of a grating will cancel, resulting in no response from the RGC (Supplementary Figure 2B). However, this cancellation does not occur with nonlinear subunits, and the RGC generates distinctive responses at twice the grating modulation frequency (‘F2’ responses, Figure 4A-B, Supplementary Figure 2B).

**Figure 4.**
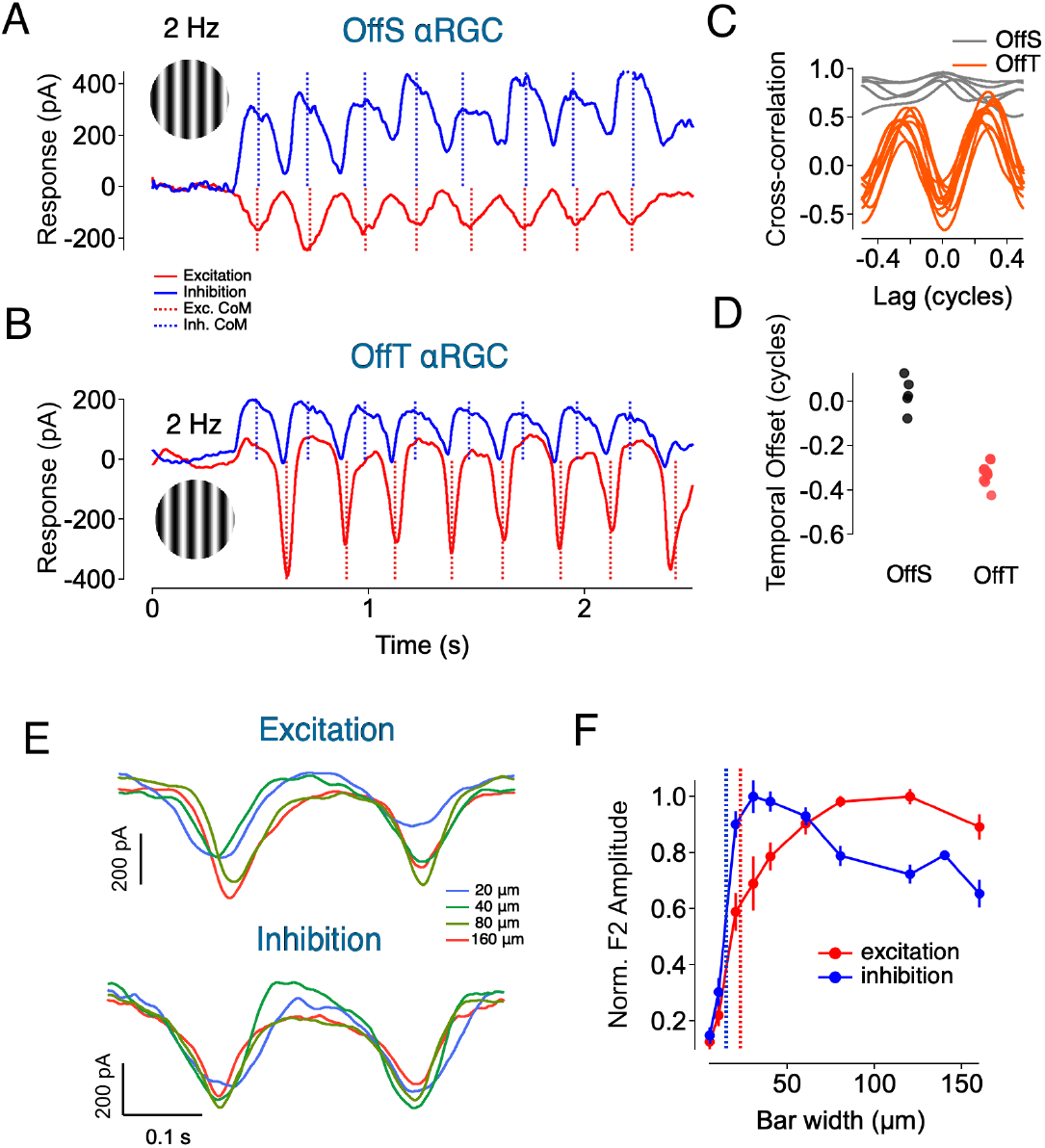
Subunits in synaptic inputs of αRGCs. (A-B) Overlay of excitatory and inhibitory currents recorded from OffS (A) and OffT (B) αRGCs at bar size of 40 μm. Dotted lines indicate centers of mass (CoM) for each cycle. (B) Cross-correlation between excitatory and inhibitory inputs for OffS and OffT αRGCs. Lag time is normalized to the F2 response cycle of 500ms. (C) Temporal offset between excitatory and inhibitory inputs for OffS and OffT αRGCs, calculated using the center of mass (CoM) of the responses. Temporal offset time is normalized to the F2 response cycle of 500ms. (D) Example excitatory (Upper) and inhibitory (Lower) currents recorded in response to contrast-reversing gratings of different bar widths. (E) Normalized F2 amplitude of excitatory and inhibitory inputs as a function of bar width. Dashed line indicates exc (red) and inh (blue) subunit radius respectively. Inhibitory inputs show a decline in amplitude for larger bar widths, while excitatory inputs plateau.

Gratings elicited excitatory and inhibitory inputs to OffS and OffT cells with distinct temporal relationships. In OffS αRGCs, excitatory and inhibitory synaptic inputs increased and decreased together with minimal temporal offset. In OffT αRGCs, inhibitory input preceded excitatory input by nearly half a grating cycle (Figure 4A-D). This temporal offset is consistent with a circuit in which A2 amacrine cells provide both direct inhibition to the ganglion cell and presynaptic inhibition at Off bipolar terminals. Presynaptic inhibition appears to suppress the expected increase in excitatory synaptic output, which instead occurs only when presynaptic inhibition subsides. This temporal offset alone does not fully account for the strong rebound excitatory synaptic input to OffT αRGCs evident when structured stimuli are removed (Figure 3B). We will examine this component of the temporal response in greater detail in the second half of the paper.

Subunit sizes, inferred from the dependence of the response on grating bar width (Figure 4 E-F), were substantially smaller than the overall RF center size (Supplementary Figure 2). Notably, excitatory and inhibitory inputs showed distinct spatial tuning profiles: inhibitory inputs peaked at smaller bar widths than excitatory inputs and as bar width increased inhibitory inputs declined while excitatory inputs remained relatively constant (Figure 4F).

**Supplementary Figure 2.**
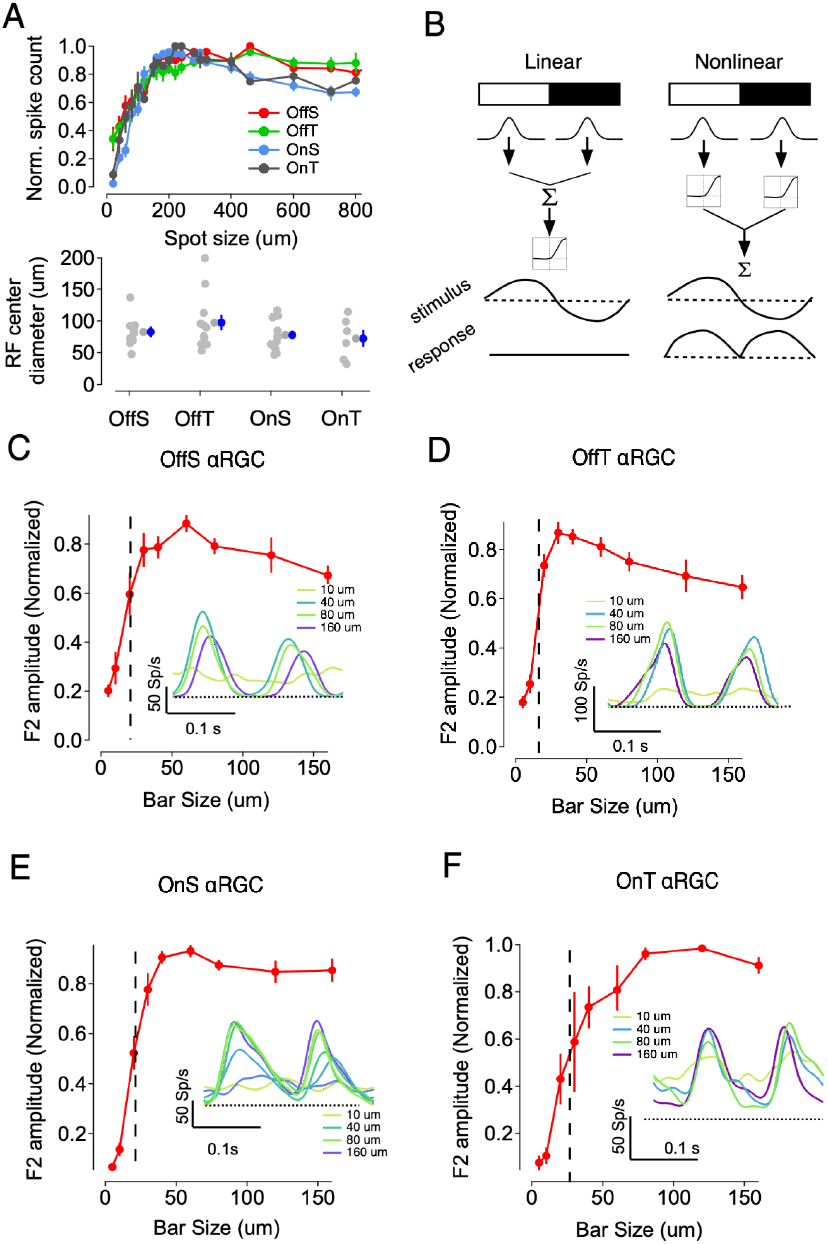
Nonlinear subunits in αRGCs receptive field. (A) Top: Area summation curves for four alpha retinal ganglion cell (αRGC) subtypes: Off-sustained (OffS), Off-transient (OffT), On-sustained (OnS), and On-transient (OnT). The summation curves plot the normalized response amplitude as a function of stimulus diameter Bottom: Scatter plots show the distribution of RF sizes. Error bars represent mean ± SEM. (B) Schematic illustration of the contrast-reversing grating stimulus used to probe the presence of nonlinear subunits in αRGC receptive fields. Nonlinear subunit receptive fields produce frequency-doubled (F2) responses when presented with such gratings, while linearly integrating receptive fields do not (C-F) Population summary data showing the nonlinear F2 response amplitude as a function of bar width. Inset: cycle average responses in an example αRGC of respective subtype.

To understand the suppression of inhibition at larger spatial scales, we constructed a spatial center-surround model of inhibitory subunits (see Methods, Supplementary Figure 3). Each subunit was modeled as a difference-of-Gaussians receptive field, embodying center-surround antagonism. By adjusting the surround strength and extent, the model accurately replicated the observed decrease in inhibitory F2 responses for wider bars (Supplementary Figure 3C-E). This strong, local surround suppression narrows inhibitory spatial integration and differentiates it from the broader integration of excitatory inputs.

The different spatial profiles of inhibitory and excitatory subunits creates a spatially dependent ratio of inhibition to excitation. Fine spatial structure drives strong local inhibition, suppressing excitation and producing a high inhibition-to-excitation ratio. But for larger, uniform stimuli, inhibitory subunits experience surround suppression and thereby contribute less inhibition, allowing excitation to dominate.

**Supplementary Figure 3.**
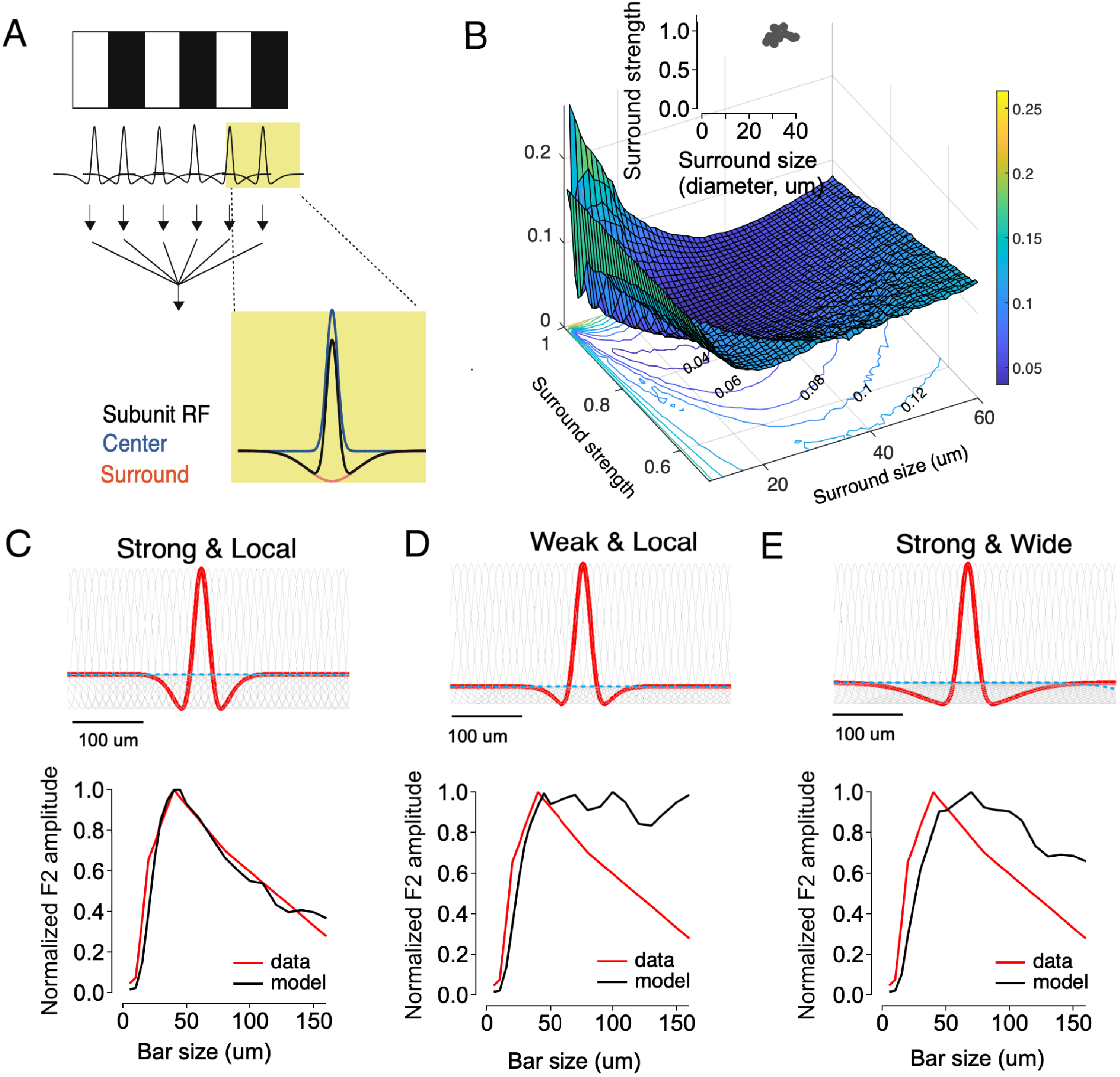
Strong and local surround mediates spatial tuning of inhibitory subunits. (A) Schematic of the center-surround subunit model for inhibitory inputs. (B) Parameter space showing the effects of surround size and strength on model fit quality. (C-E) Model fits (black) to experimental data (red) for inhibitory inputs with different surround properties: strong & local (C, surround size 15 μm, surround strength 0.9), weak & local (D surround size 15 μm, surround strength 0.2), and strong & nonlocal (E, surround size 40 μm, surround strength 0.9).

### Spatially Local Inhibition Arises from Inter-Pathway Glycinergic Modulation by A2 Amacrine Cells

Figures 1 and 2 show that OffT αRGCs exhibit a robust preference for spatially homogeneous stimuli and suggest that preference is driven by presynaptically acting inhibition that suppresses excitatory responses to fine spatial structure. To identify the specific circuit elements mediating this inhibition, we examined the interactions between the parallel rod pathways converging onto Off cone bipolar cells. The primary rod pathway involving A2 amacrine cells was of particular interest, as these neurons can provide both direct inhibition to OffT αRGCs and presynaptic inhibition onto Off cone bipolar terminals. To test whether this pathway is necessary for homogeneity preference, we selectively blocked it using two complementary approaches.

First, we disrupted the rod-to-rod bipolar synapse using a cocktail of agonists and antagonists (LY341495 and APB) of the mGluR6 glutamate receptors expressed by On bipolar cells (Figure 5A-C) (Grimes et al., 2014). Using such a cocktail prevents aberrant signaling associated with strongly hyperpolarizing bipolar cells with an agonist alone (Grimes et al., 2014). Suppressing light-dependent rod bipolar input to AII amacrine cells in this way revealed robust excitatory responses to spatially structured stimuli at onset—responses that are normally absent (Figure 5A–C). Similarly, application of strychnine to block glycinergic transmission—the inhibitory transmitter released by A2 cells (Wässle et al., 2009)—restored strong onset excitation to spatial structure (Figure 5E–F). Both manipulations also attenuated the robust offset responses normally observed, supporting the role of presynaptic inhibition in regulating synaptic gain across stimulus transitions.

**Figure 5.**
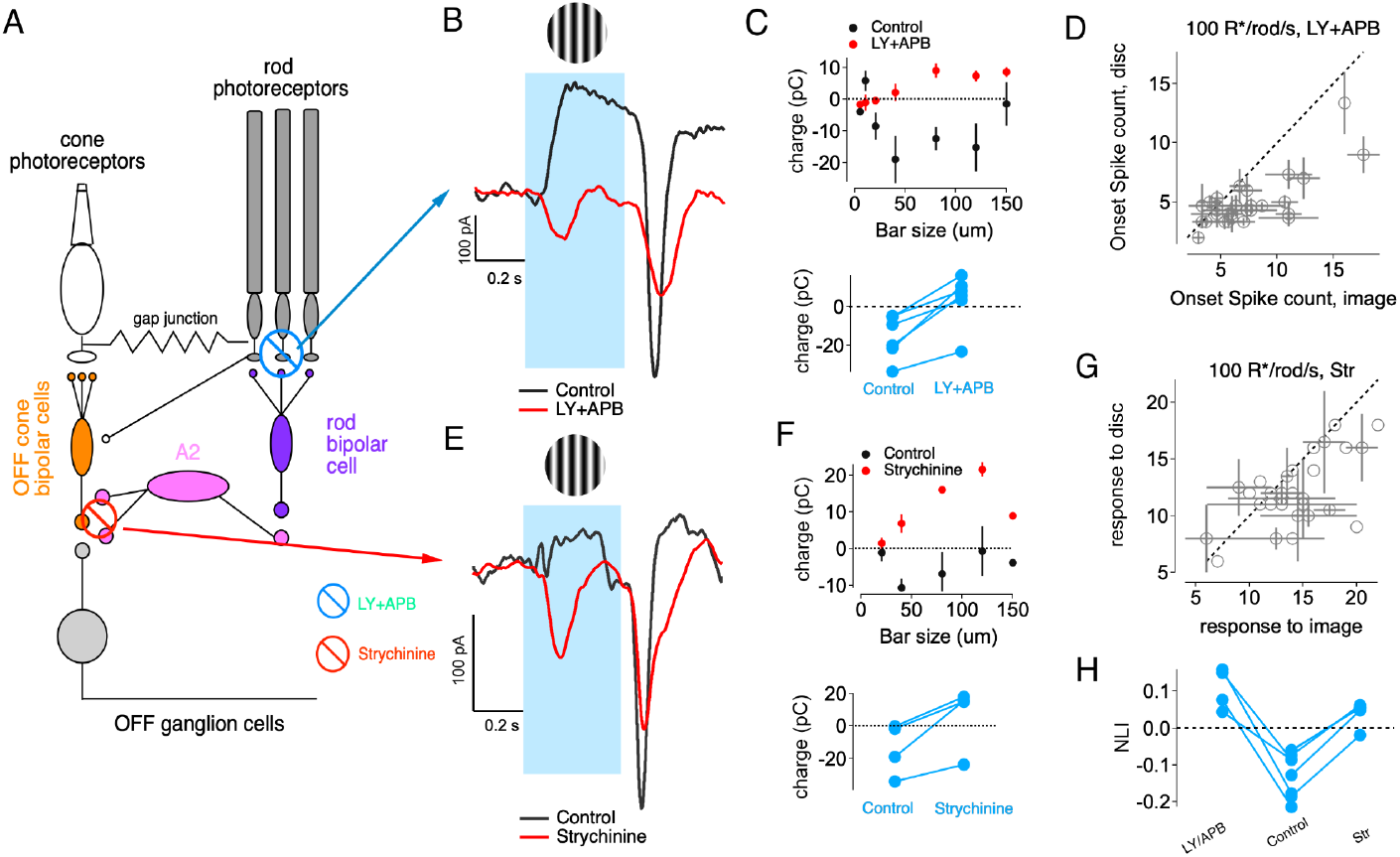
Pharmacological profiles of presynaptic inhibition in OffT αRGCs. (A) Schematic showing parallel rod pathways in the retina with sites of pharmacological intervention. LY341495+APB (blue) blocks the rod->rod bipolar cell transmission, while strychnine (red) blocks glycinergic transmission from A2 amacrine cells. (B) Representative excitatory current traces in response to flashed gratings under control conditions (black) and with LY341495 (10 μM) +APB (10 μM) application (red) to block the primary rod pathway. Blue shading indicates grating presentation. (C) Top: Quantification of excitatory charge transfer in response to onset of gratings with different bar sizes under control (black) and LY341495+APB (red) conditions. Bottom: Paired comparison of excitatory charge transfer for individual cells under control and LY341495+APB conditions. (D) Comparison of OffT αRGC onset spike responses to natural image patches (x-axis) versus linear-equivalent discs (y-axis) under LY341495+APB treatment at 100 R*/rod/s. (E) Representative excitatory current traces in response to flashed gratings under control conditions (black) and with strychnine (0.5 μM) application (red) to block glycinergic transmission. (F) Top: Quantification of excitatory charge transfer in response to gratings with different bar sizes under control (black) and strychnine (red) conditions. Bottom: Paired comparison of excitatory charge transfer for individual cells under control and strychnine conditions. (G) Comparison of OffT αRGC onset spike responses to natural image patches versus linear-equivalent discs under strychnine treatment at 100 R*/rod/s. (H) Summary of nonlinearity indices (NLI) for onset spike responses under control conditions, LY341495+APB treatment, and strychnine treatment at 100 R*/rod/s. Each line represents an individual cell.

When tested with natural images, both pharmacological interventions abolished the homogeneity preference, inverting the normal response pattern such that OffT αRGCs now preferred spatially structured images over linear-equivalent discs (Figure 5D,G-H). This confirms that the homogeneity preference requires A2-mediated inhibition rather than being an intrinsic property of the excitatory pathway.

Collectively, these findings support a model in which the homogeneity preference in OffT αRGCs arises from the interaction between parallel pathways, rather than from mechanisms confined to the primary pathway. The key player in this interaction appears to be A2 amacrine cells, which provide both presynaptic inhibition to Off cone bipolar cells and direct inhibition to OffT αRGCs. This dual inhibitory action effectively suppressed excitatory signals from secondary and tertiary pathways in response to spatial structure, thereby shaping the unique response properties of OffT αRGCs to natural images.

### Light-Level Dependent Shifts in OffT αRGC Spatial Computation

Given that A2-mediated inhibition via the primary rod pathway is required to generate the homogeneity preference in OffT αRGCs, we next asked how this computation changes with light adaptation. As ambient light levels increase, the primary rod pathway becomes progressively suppressed, and A2 amacrine cells are instead driven predominantly through electrical coupling with ON cone bipolar cells (Ke et al., 2014; Grimes et al., 2018). This shift in input source is expected to alter the strength and spatial organization of A2-mediated inhibition. To examine the functional consequences of this circuit reconfiguration, we first assessed whether the balance of excitatory and inhibitory synaptic inputs to OffT αRGCs changes with light level. At 1000 R*/rod/s—now within the photopic range—compared to 100 R*/rod/s in the mesopic range, we observed a clear reduction in inhibitory conductances relative to excitatory conductances (Figure 6A–C). This shift in E/I balance reflects the attenuation of A2-driven inhibitory input, consistent with reduced engagement of the primary rod pathway at higher light levels, and an increase in excitatory input. This shift in E/I balance produced a reversal of spatial sensitivity: after adaptation to photopic conditions, OffT αRGCs responded more strongly to spatially structured image patches than to homogeneous discs—opposite the pattern observed under mesopic conditions (Figure 6D–F).

**Figure 6:**
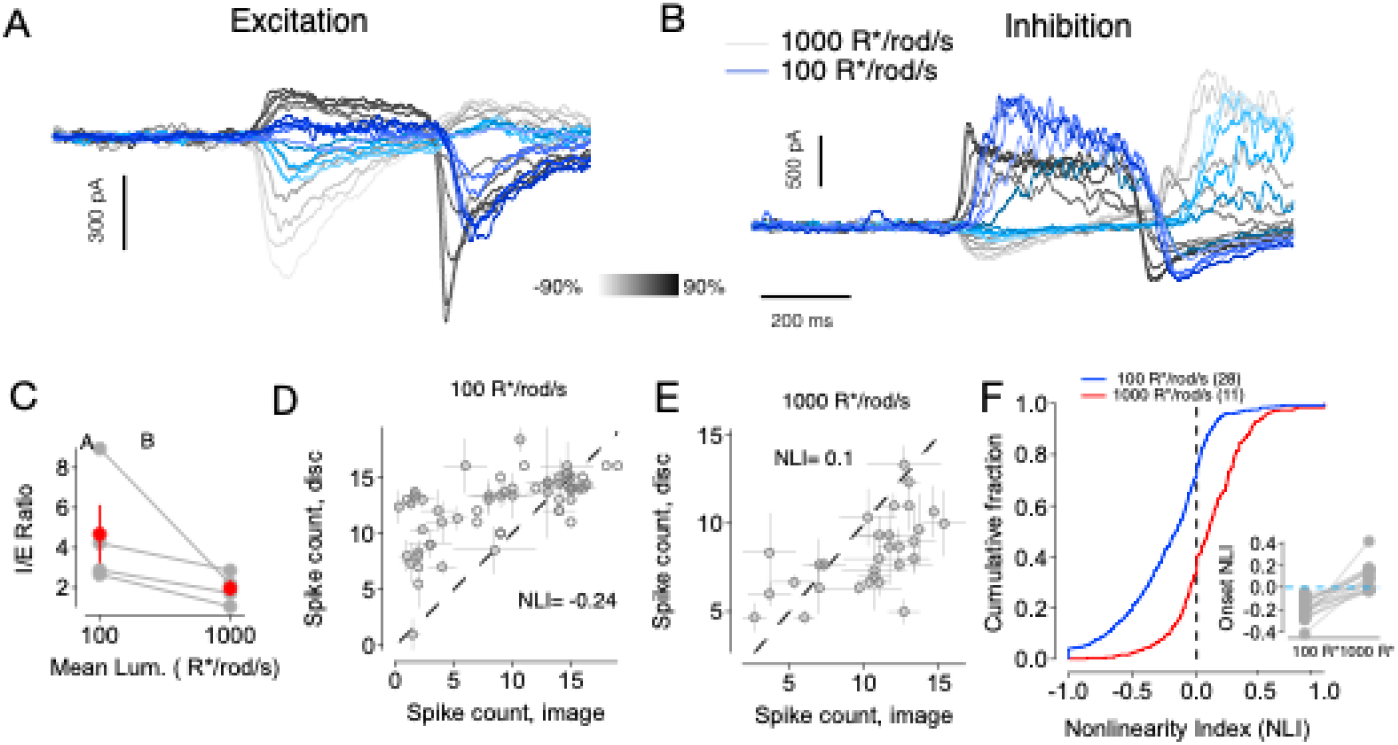
Luminance dependent spatial encoding in OffT αRGCs. (A) Excitatory currents recorded from an example OffT αRGC in response to spot stimuli at two mean light levels across various contrast steps. Red traces: 1000 R*/rod/s, blue traces: 100 R*/rod/s. (B) Inhibitory currents recorded from the same cell under the same conditions as in (A). (C) I/E ratio for individual cells (gray) and population average (red) at the two light levels. (D) Spike count responses of an example OffT αRGC to natural image patches versus linear equivalent discs at 100 R*/rod/s. Each point represents one image patch. (E) Responses of the same cell at 1000 R*/rod/s. (F) Cumulative distributions of Nonlinearity Index (NLI) values across image/disc pairs and population of OffT αRGCs at 100 R*/rod/s (yellow, n=28 cells) and 1000 R*/rod/s (red, n=11 cells).

These results demonstrate that light-level dependent shifts in rod pathway routing alter both the strength of A2-mediated inhibition and the spatial computations performed by OffT αRGCs, highlighting the essential role of A2-mediated pathway in dynamically shaping spatial computations in the retina.

### Short-Term Plasticity and Glycinergic Inhibition Underlie History-Dependent Excitation in OffT αRGCs

We next investigated the robust rebound of excitatory input observed at the offset of spatially structured stimuli (e.g. Figure 2A, 3B). We hypothesized that this rebound reflects a history-dependent modulation of synaptic gain, mediated by glycinergic inhibition and short-term plasticity at the bipolar terminal. This idea is consistent with previous work (Ke et al., 2014), which showed that brief periods of reduced drive potentiate subsequent responses via recovery from vesicle depletion or disinhibition at bipolar outputs.

To test this hypothesis, we adapted a paired-pulse paradigm in which two identical dark flashes (−90% contrast, 0.3s) were separated by an intervening contrast step presented on a 100 R*/rod/s background. The contrast of the intervening step ranged from 0 (return to baseline) to positive values (opposing), and we measured both synaptic input and spiking output of OffT αRGCs across conditions (Figure 7A–D). For both zero-contrast and positive-contrast intervals, the response to the second pulse was substantially larger than that to the first pulse, often as much as 3-fold. During the period between the pulses both firing rate and excitatory synaptic input were substantially reduced. This suggests that brief periods of reduced drive potentiate subsequent responses, likely through recovery from synaptic depression. The strength of this facilitation depended on the duration of the recovery interval, with the paired-pulse ratio decreasing progressively with longer intervals (200ms to 5000ms) before plateauing (Figure 7E-H). We also tracked baseline activity (measured immediately before each second stimulus pulse) to monitor the dynamics of tonic release. The baseline gradually returned to pre-stimulus levels over a similar timescale (Figure 7H).

**Figure 7:**
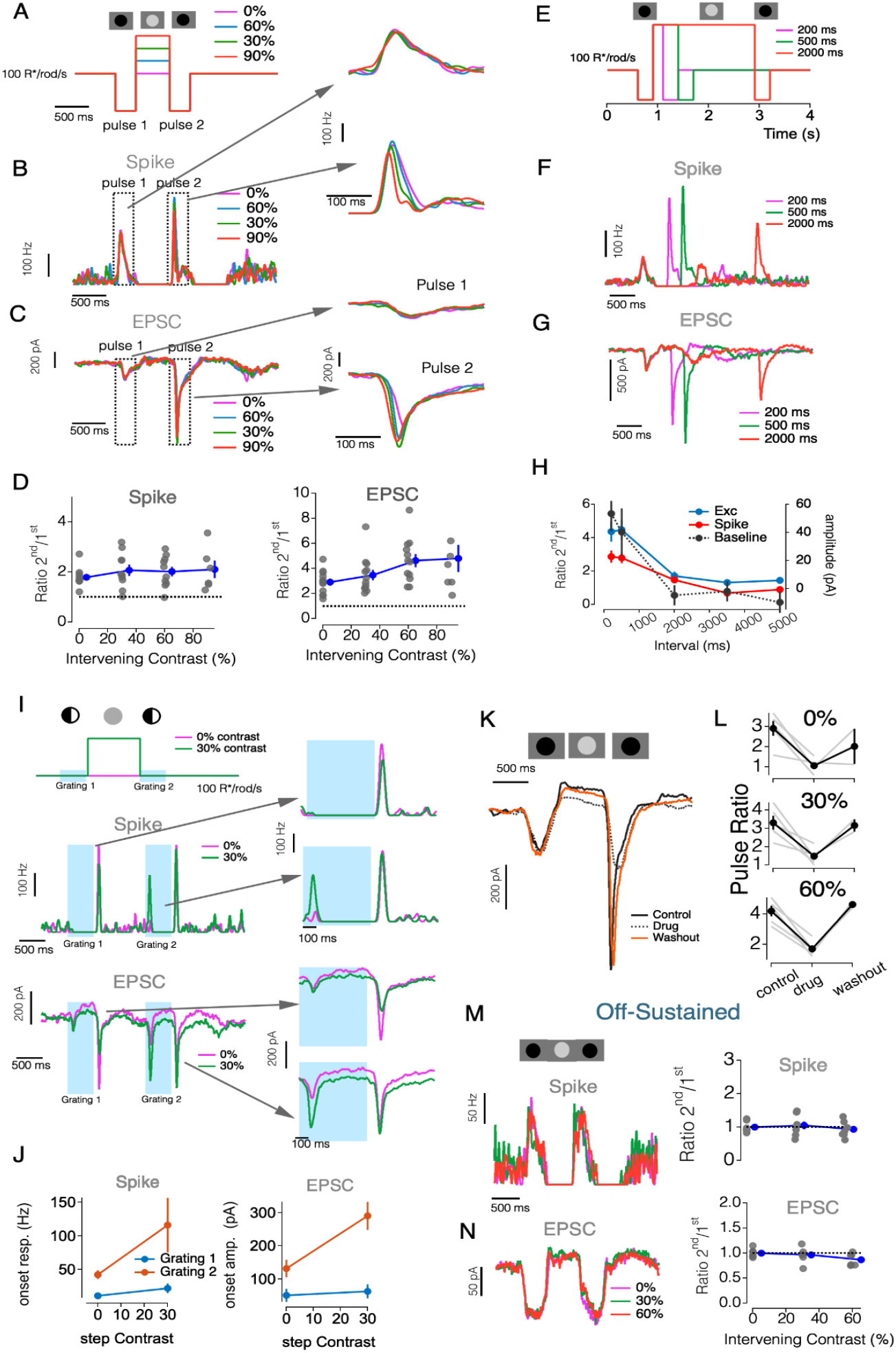
Temporal Dynamics of Synaptic Interactions in Off αRGCs. (A-H). Bright Interstimulus Intervals Enhance Subsequent Light Responses in OffT αRGCs: (A) Stimulus protocol showing two negative light pulses (300 ms, −90% contrast) separated by various intervening contrasts (0%-90% contrast, 500 ms). (B) Spike responses of an OffT RGC to paired pulses. Insets: expanded views of responses to pulse 1 (top) and pulse 2 (bottom), aligned to pulse onset. (C) Excitatory postsynaptic currents (EPSCs) evoked by paired pulses in the same cell as (B). (D) Spike (Left) and EPSC (Right) paired-pulse ratio (pulse 2/pulse 1 peak amplitude) as a function of interpulse contrast. Error bars, ±SEM across cells. (E) Stimulus protocol showing two negative light pulses (300 ms, −90% contrast) separated by variable duration intervals (200 ms, 500 ms, and 2000 ms; 60% contrast). (F) Spike responses of an OffT αRGC to paired pulses separated by different intervals. Traces show responses for 200 ms (magenta), 500 ms (green), and 2000 ms (red) intervals. (G) Excitatory postsynaptic currents (EPSCs) evoked by paired pulses in the same cell as (F). (H) Spike (blue), EPSC (red), and baseline (black) paired-pulse ratios as a function of interpulse interval duration. Error bars, ±SEM across cells. (I) Upper: Stimulus protocol showing two sequential grating stimuli (500 ms, 90% spatial contrast) separated by various intervening contrast steps (0%-30% contrast, 1000 ms). Middle:spike responses to two sequential grating stimuli (grating 1 and grating 2, 500 ms each). Insets: expanded views of responses to grating 1 and grating 2. Lower: EPSCs evoked by sequential grating stimuli in the same cell as (F). (J) Spike (H) and EPSC (I) responses amplitudes to grating 1 and grating 2 as a function of inter-grating contrast (0-30%). Error bars, ±SEM across cells. (K-L) Glycinergic inhibition is necessary for relief from suppression in OffT αRGCs: (K) Example excitatory current traces in response to paired pulses under control conditions (solid black line), with strychnine application (0.5μM, dotted black line), and after washout (orange line). (L) Summary of paired-pulse ratios under control, strychnine, and washout conditions for different intervening contrasts (0%, 30%, and 60%). Gray lines show individual cells, black lines with error bars show mean ±SEM. (M-N) OffS αRGCs show minimal relief from suppression effects: (M) Example spike (left) responses of an OffS αRGC to paired pulse stimuli with different intervening contrasts (0%, 30%, and 60%). Right: paired-pulse ratios for spike responses in OffS αRGCs as a function of intervening contrast. Gray dots show individual cells, and blue line with error bars show mean ±SEM. (N) same as (M) but for EPSC in OffS αRGC.

We next asked whether the same mechanism shapes responses to spatially structured stimuli by presenting two successive high-contrast gratings (Figures 7I-J). Again, a brighter interval separating the flashed gratings elicited a larger response to onset of the second grating in both spike and EPSC measurements (Figure 7I-J). To determine whether glycinergic inhibition is necessary for this facilitation, we repeated the paired-pulse experiments in the presence of strychnine (Figures 7 K-L). Suppressing glycinergic transmission eliminated the second-pulse enhancement, confirming that presynaptic inhibition—most likely via A2 amacrine cells—leads to response facilitation.

Finally, we asked whether OffS αRGCs show a similar history-dependent increase in excitatory gain (Figures 7M-N). Even with high intervening brightness, OffS cells exhibited little or no facilitation. This lack of facilitation occurs even though the firing rate and tonic excitatory input between the two pulses were strongly suppressed. These results underscore that the interaction between glycinergic inhibition and short-term synaptic plasticity is a distinctive feature of OffT αRGCs.

Taken together, these experiments identify a dual role for inhibition in shaping OffT αRGC activity. First, it suppresses excitatory drive during structured or dark stimuli. Second, by allowing bipolar terminals to recover from depression, it establishes a powerful rebound of excitatory output once inhibition lifts. This mechanism endows OffT, but not OffS, αRGCs with flexible, context-dependent control over their excitatory gain.

### A Unified Spatial-Temporal Model Reproduces OffT αRGC Homogeneity Preference

Our analyses identified two key mechanisms underlying homogeneity preference in OffT αRGCs: (1) a spatial mismatch between narrowly-tuned inhibitory inputs and broadly integrating excitatory inputs, and (2) presynaptic inhibition at bipolar terminals that enables recovery from chronic synaptic depression and a corresponding large increase in excitatory input at the offset of spatially-structured images. We next sought to develop a unified model that integrates these complementary mechanisms (Figure 8).

**Figure 8.**
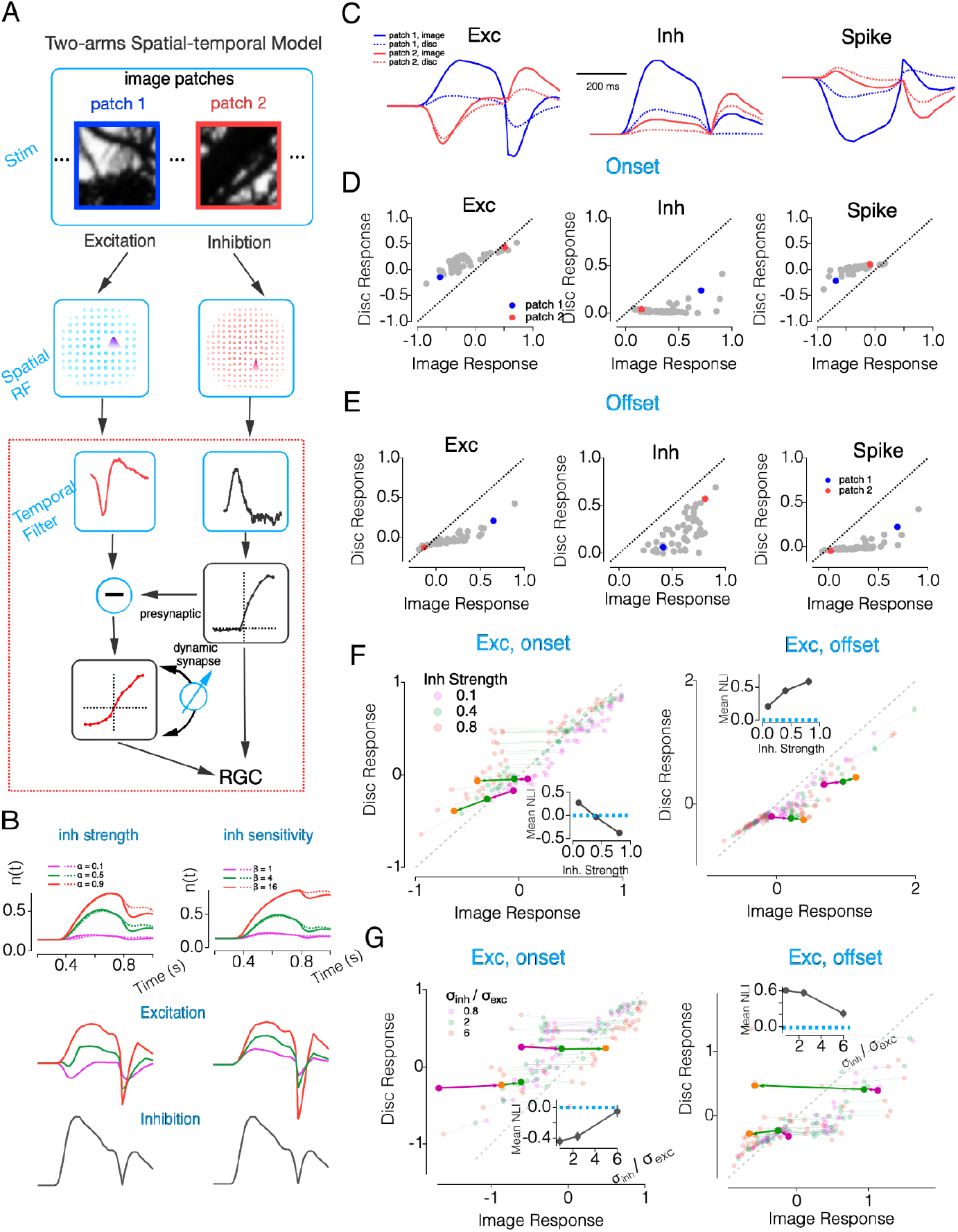
Spatial-Temporal Dynamics of Excitation/Inhibition Balance Mediate Homogeneity Preference in the Retina. (A) Schematic of the two-stream spatial-temporal model. Image patches (top) are processed through parallel excitatory and inhibitory pathways with distinct spatial receptive fields (middle row). Both streams then pass through temporal filters. The inhibitory pathway modulates the excitatory pathway through presynaptic inhibition, which operates via a dynamic synapse with depression (blue feedback arrow). The final output integrates these pathways to form the retinal ganglion cell response. (B) Dynamic simulations of the temporal-only model (dashed rectangle in A) showing vesicle occupancy rate and excitatory output under different parameter conditions in response to a flashed grating (as in Figure 3). Left panel: Vesicle occupancy dynamics (n(t)) with varying presynaptic inhibition strengths (α = 0.1, 0.4, 0.8). Right panel: Vesicle occupancy dynamics with varying inhibition sensitivity values (β = 1, 4, 8). Corresponding excitatory outputs and inhibitory traces are shown below each condition. (C) Example conductance response traces showing excitatory conductance trace (left), inhibitory conductance trace (middle), and spike responses (3Exc-Inh, right) to two natural image patches (solid lines) and their equivalent homogeneous discs (dotted lines). The spatial structure in patch 1 (blue) and patch 2 (red) evokes distinct patterns of excitation and inhibition. (D) Comparison of onset responses to natural image patches versus their equivalent discs for excitation (left), inhibition (middle), and spike output (right). Inhibition strongly prefers structured images (points below unity line) while excitation prefers homogeneous discs (points above unity line). (E) Similar to C, but showing offset responses after stimulus termination. (F) Simulation results showing how inhibitory strength modulates spatial integration. Scatter plots compare image versus disc responses at stimulus onset (left) and offset (right) with different inhibitory strengths (0.1, 0.4, 0.8). Responses represent area sums (unitless) normalized to the maximal response during stimulus onset across all conditions. Semi-transparent points represent individual image patches, while thick colored lines show responses for a few example patches. Insets show how the mean nonlinearity index (NLI) changes with inhibitory strength. (G) Same as F, but for inhibitory/excitatory subunit size ratio (0.8, 2, 4). Insets show how the mean NLI changes with the subunit size ratio.

**Supplementary Figure 4.**
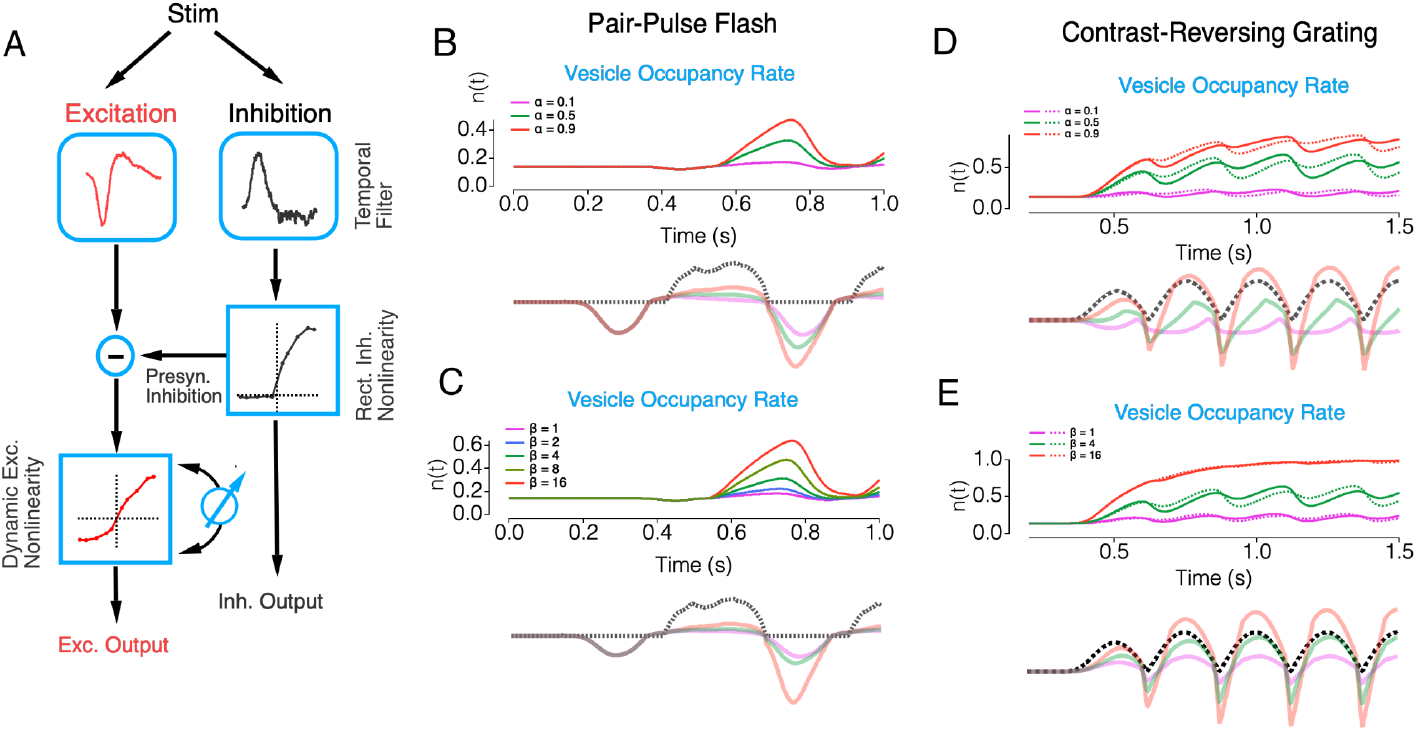
Dynamic Synaptic Depression Mediates History-Dependent Excitation/Inhibition Interaction in Retinal Processing. (A) Schematic of the temporal dynamic synapse model showing the excitatory pathway (red) and inhibitory pathway (black). Inhibition modulates excitation through presynaptic inhibition, with inhibitory pathways passing through rectifying nonlinearity while excitatory pathways passing through piecewise nonlinearity. The blue arrow indicates the dynamic regulation of excitatory output through vesicle occupancy. (B-C) Simulated responses to paired-pulse flash stimuli. Top panels show vesicle occupancy rate (n(t)) and bottom panels show excitatory conductance output (colored lines) along with inhibitory current (dashed black line). (B) demonstrates the effect of varying presynaptic inhibition strength (α) while (C) shows responses with different inhibition sensitivity values (β). (D-E) Model responses to contrast-reversing grating stimuli.

**Supplementary Figure 5:**
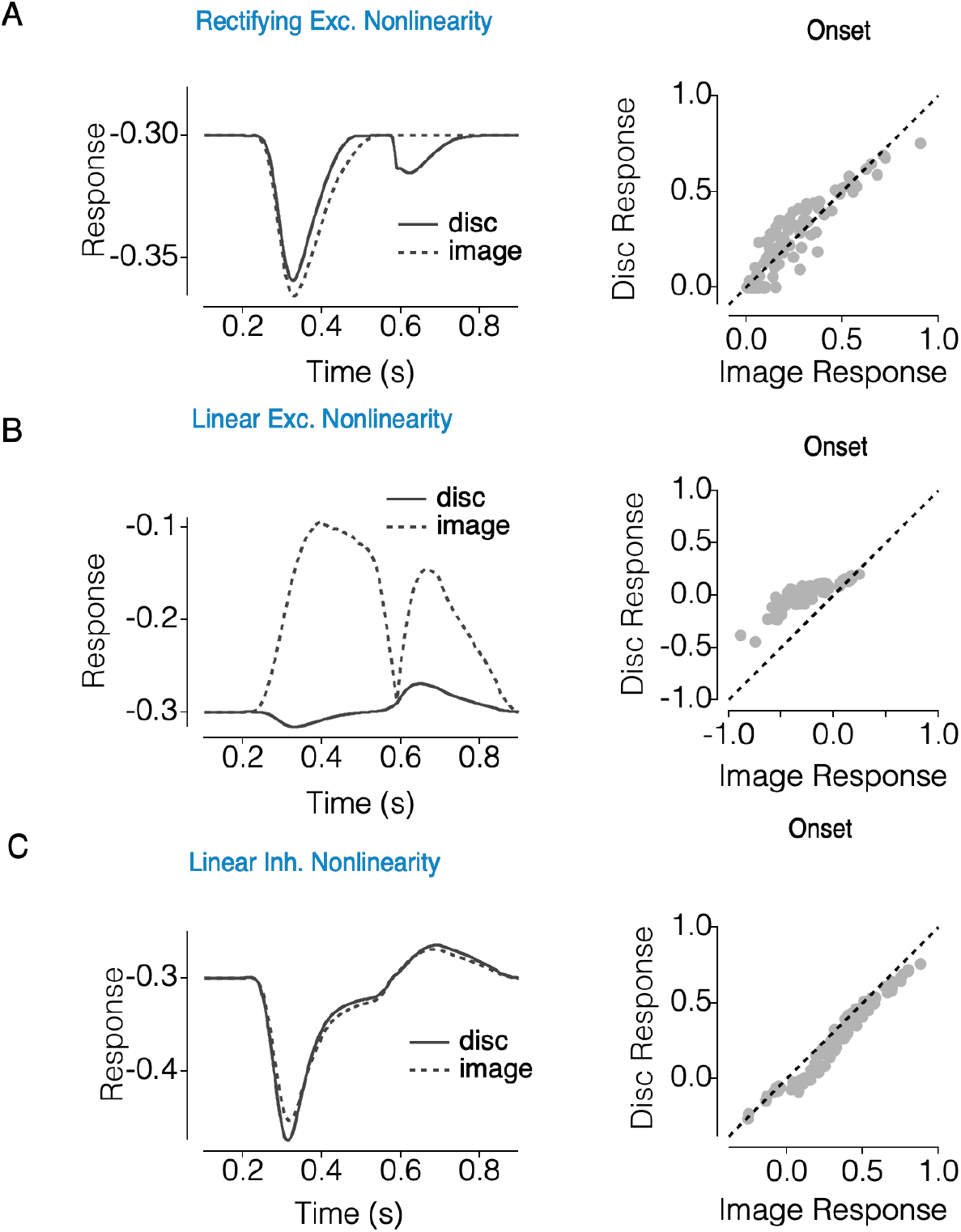
Effects of excitatory and inhibitory nonlinearities on spatial integration in the retina. (A) Example response of the model with rectifying excitatory nonlinearity to natural image patches (dashed line) and their corresponding linear equivalent discs (solid line). Left panel shows example response traces over time. Right panel shows the scatter plot of image responses versus disc responses at stimulus onset, with most points clustered along the unity line (dashed), indicating similar responses to both stimulus types. (B) Response of the model with linear excitatory nonlinearity to the same stimuli. Left panel shows example traces demonstrating suppression of responses to natural images (dashed line) compared to disc stimuli (solid line). Right panel shows the scatter plot of responses at stimulus onset. (C) Response of the model with linear inhibitory nonlinearity. Left panel shows example traces with similar responses to both natural images (dashed line) and linear equivalent discs (solid line). Right panel shows the scatter plot of responses at stimulus onset, with points clustered along the unity line, indicating that linear inhibition does not generate differential responses to spatial structure.

Our model builds on a hybrid LN framework (Figure 8A): the excitatory pathway is a dynamic LN model, in which the slope of the output nonlinearity—reflecting synaptic gain—is multiplicatively scaled by the vesicle occupancy (n), which evolves according to a depletion-and-recovery process. This allows presynaptic inhibition to directly modulate gain by reducing vesicle release rates (see Methods). The inhibitory pathway is implemented as a conventional static LN model, without vesicle dynamics.

We started with a reduced temporal-only implementation (Figure 8B, Supplementary Figure 4A, also see Methods) that focuses on core temporal dynamics. This simplified temporal model tracks excitatory and inhibitory responses across two visual hemifields receiving either matching or opposing contrast inputs, allowing us to isolate temporal mechanisms in responses to a flashed grating (as in Figure 3) in a more constrained parameter space before extending to the full spatial-temporal domain. The two-hemifield model successfully replicates two characteristic features of OffT αRGC responses to a flashed grating: suppression of excitatory conductance during stimulus onset followed by rebound excitation at offset (Figure 8B). These effects emerge directly from the presynaptic suppression of vesicle release during stimulation, followed by enhanced release once inhibition is relieved at grating offset. Both the suppression and rebound scale predictably with the strength of presynaptic inhibition strength (*α*) and its sensitivity parameter (*β*). A higher β increases the coupling between inhibition and vesicle dynamics, making synaptic output more sensitive to small changes in inhibitory input (Figure 8B; see Methods). Our model also captures other temporal signatures of OffT αRGCs in response to additional stimuli, including paired-pulse facilitation and phase-shifted responses to contrast-reversing gratings (see Supplementary Figure 4).

The full spatial-temporal model (Figure 8A) adds the distinct spatial properties of excitatory and inhibitory subunits observed experimentally. This model successfully reproduced both suppression of onset responses and enhancement of offset responses to structured natural images (Figure 8C-E). These effects emerged from spatially restricted inhibition suppressing excitatory drive to structured stimuli and permitting recovery of excitatory synaptic gain. Consistent with our pharmacological experiments (Figure 5), reducing inhibition attenuated the predicted homogeneity preference (Figure 8F). Increasing inhibitory subunit size similarly reduced the model’s preference for homogeneous stimuli (Figure 8G). Replicating the observed homogeneity preference also depended on nonlinear properties of both excitation and inhibition (Supplementary Figure 5).

This spatial-temporal framework captures the core computations shaping OffT αRGC responses: structured stimuli activate inhibitory subunits that suppress excitation and permit recovery of synaptic gain, while transitions to homogeneity reveal enhanced excitation through disinhibition and gain facilitation. This model provides a mechanistic basis for understanding how nonlinear spatial integration and history-dependent gain interact to shape retinal output under naturalistic conditions.

## Discussion

OffT αRGCs exhibit unexpected responses to spatial stimuli - responding more strongly to homogeneous stimuli than to those with spatial structure. This is unlike the responses of most RGCs, which are enhanced by spatial structure due to nonlinear receptive field subunits. The unusual spatial selectivity of the OffT αRGCs suggests that mechanisms not captured by the canonical view of retinal processing shape their responses. Here, we identified two circuit mechanics that together can account for OffT spatial selectivity. First, synaptic inhibition was tuned to finer spatial scales than excitation, causing a preferential cancelation of responses to fine spatial structure. Second, the excitatory synapses made onto OffT αRGCs exhibited strong short-term synaptic plasticity and the resulting modulation of synaptic gain by presynaptic inhibition amplified responses at the offset of spatial structured stimuli.

### Non-canonical roles of inhibition in spatial integration

The stimulus selectivity of many sensory neurons is shaped by interactions between excitation and inhibition. A canonical form of such interactions is lateral inhibition - in which inhibition is tuned to a broader set of stimuli than excitation and hence can sharpen tuning by suppressing responses to non-optimal stimuli (Pouille et al., 2009; Shapley and Xing, 2013). Another canonical form is feedforward inhibition, in which an extra synapse in the inhibitory pathway causes a short temporal delay between excitation and inhibition and a corresponding brief time window in which excitation can dominate the response (Murphy and Rieke, 2006; Ferrante et al., 2009; Keck et al., 2012). Both of these forms of inhibition are prominent in retinal circuits (Diamond, n.d.).

Inhibition can also contribute to computation in non-canonical ways. Suppressing inhibition eliminated characteristic properties of the responses of homogeneity detectors in salamander retina (Bölinger and Gollisch, 2012). This suggests a role very different from classic lateral inhibition (Barlow, 1953). Indeed, we found that inhibitory input to OffT RGCs was tuned to finer spatial scales than excitatory input. This difference broadens spatial tuning by allowing inhibition to suppress responses to fine but not coarse spatial structure. Excitation and inhibition typically share many circuit elements, and hence the stimulus selectivity of inhibition inherits many features from excitation (Anderson et al., 2000; Monier et al., 2003; Priebe and Ferster, 2005; Cardin et al., 2007; Denève and Machens, 2016). The more narrowly tuned inhibition that we observe here requires instead that inhibition originates from circuitry that is largely distinct from excitation, and indeed the preference of OffT RGCs for homogeneous inputs was strongest when excitation and inhibition originated through parallel and distinct retinal circuits – excitation through cone bipolar circuits and inhibition through rod bipolar circuits. The key aspect of this parallel processing is that it allows for differential tuning of excitation and inhibition.

### Synaptic Depression as Mechanism for Context-Dependent Processing

History-dependent changes in synaptic strength shape responses in many circuits. For example, depression at excitatory synapses often serves as a gain control mechanism (Ke et al., 2014) whereas depression at inhibitory synapses, and the resulting disinhibition, can produce facilitation (Kastner et al., 2019; Heintz et al., 2022). These effects occur through a variety of synaptic mechanisms and correspondingly operate on a variety of timescales. Retinal signals are strongly shaped by short-term plasticity due to vesicle depletion and the resulting modulation of synaptic gain (von Gersdorff and Matthews, 1997; Rabl et al., 2006; Singer and Diamond, 2006; Oesch and Diamond, 2011).

Our findings expand upon this framework, illustrating how synaptic depression at bipolar terminals, when combined with spatially precise glycinergic inhibition, generates a novel form of spatial selectivity. In the absence of a stimulus, we find that the excitatory synapses onto OffT RGCs are tonically active, and correspondingly operate at low gain. Suppression of this tonic activity allows recovery of synaptic gain and large responses to subsequent stimuli. Presynaptic inhibition is a key source of such suppression, and this contributes to the homogeneity preference of these cells. Specifically, the onset of spatially structured stimuli elicit strong presynaptic inhibition which suppresses vesicle release. This allows vesicle replenishment and enhances the excitatory responses generated at a transition from spatially-structured to homogeneous stimuli.

Interestingly, excitatory inputs to OffS cells exhibit a similar level of tonic activity but do not show a recovery of gain following suppression of this tonic activity. Thus different properties of different bipolar synapses shape responses and contribute to diversification of RGC responses. A similar difference in short-term plasticity at different bipolar cell synapses has been described for On transient versus On sustained retinal ganglion cells (Kuo et al., 2024). On transient cells exhibit more pronounced synaptic depression—consistent with smaller or more rapidly depleting vesicle pools at presynaptic bipolar terminals—whereas On sustained cells maintain higher baseline release with less depletion during prolonged stimuli.

### Divergence of OffS and OffT Pathways

The functional asymmetry between OffT and OffS αRGCs at mesopic light levels is striking given the similarities in the circuitry that controls their responses. Both cell types receive excitatory synaptic input from Off bipolar cells and inhibitory synaptic input primarily from A2 amacrine cells at the light levels used here. However, excitatory inputs to OffT and OffS differ in the strength of presynaptic inhibition and in short-term synaptic plasticity that modulates synaptic gain. At excitatory synapses made onto OffT cells, glycinergic inhibition from A2 amacrine cells both suppresses excitatory drive during structured or dark stimuli and sets up a strong rebound of excitation once that inhibition is relieved—an effect we do not observe in OffS cells.

Mechanistically, this disparity likely originates from distinct synaptic profiles: OffS αRGCs rely heavily on type 2 bipolar cells, with additional inputs from type 3a and type 4 bipolar cells, whereas OffT cells receive input predominantly from type 3a, 3a and 4 bipolar cells (Sabbah et al., 2018; Yu et al., 2018). A2 amacrine cells contact all of these Off-bipolar subtypes in the inner plexiform layer (Tsukamoto and Omi, 2017; Graydon et al., 2018), but the magnitude and dynamics of presynaptic inhibition—and the intrinsic release properties of each bipolar subtype—may differ substantially. Further research into the intrinsic properties of Off-cone bipolar subtypes and their synaptic interactions with A2 amacrine cells could elucidate the root cause of this asymmetry.

## Methods and Materials

### Electrophysiology

Experiments were conducted on whole-mount retinas from dark-adapted wild-type (C57/BL6 or sv-129, either sex, 1–12 months)) mice using procedures in compliance with the University of Washington’s Institutional Animal Care and Use Committee. The retina was isolated with Retinal Pigment Epithelium (RPE) and sclera attached and stored in oxygenated (95% O2/5% CO2) Ames medium (Sigma) at 32°C–34°C. Isolated retinas were flattened onto polyL-lysine slides before being transferred under the microscope for recording and were continuously perfused with Ames medium at a flow rate of approximately 8 mL/min. For cell-attached and whole-cell recordings, retinal neurons were visualized and targeted using infrared light (>950 nm).

Voltage-clamp recordings were obtained using pipettes (for RGCs, 2–3 MΩ; for bipolar cells, 10–14 MΩ) filled with an intracellular solution containing (in mM) the following: 105 Cs methanesulfonate, 10 TEA-Cl, 20 HEPES, 10 EGTA, 2 QX-314, 5 Mg-ATP, 0.5 Tris-GTP, and 0.1 Alexa-750 hydrazide (*~*280 mOsm; pH *~*7.3 with CsOH). Current clamp recordings for A2 amacrine cells were conducted with pipettes ~8 MΩ using an intracellular solution containing (in mM) the following: 123 K-aspartate, 10 KCl, 10 HEPES, 1 MgCl2, 1 CaCl2, 2 EGTA, 4 Mg-ATP, 0.5 Tris-GTP, and 0.1 Alexa-750 hydrazide (*~*280 mOsm; pH *~*7.2 with KOH). NBQX (10 μm; Tocris) or TTX (0.5 μm; Alamone) was added to the perfusion solution. Recording for rod bipolar cells was kept brief (typically <5 min) to reduce washout effects. Absolute voltage values were not corrected for liquid junction potentials (*K* ^+^ based = −10.8 mV; *Cs*^+^ based = −8.5 mV).

### Visual Stimuli

Stimuli were generated, and visual response data was acquired using custom-written stimulation and acquisition software packages Stage (http://stage-vss.github.io) and Symphony (http://symphony-das.github.io), respectively. These stimuli were displayed at a frame rate of 60 Hz on an OLED microdisplay monitor (800 x 600 pixels, eMagin), with a focus through a 10x objective, directly onto the photoreceptors. The configuration resulted in a resolution of either 1.8 μm/pixel or 1 μm/pixel at the retina, respectively, for two different recording rigs used. Stimuli were calibrated using the measured monitor optical power output, the spectral content of the monitor, mouse photoreceptor spectral sensitivity, and a collecting area of 0.5 μ*m*^2^ for mouse rods. Unless otherwise specified, mean light levels produced *~*100 isomerizations per rod per second (R*/rod/s).

At the beginning of each recording, the cell’s receptive field center was identified using a split-field contrast reversing grating stimulus at 2Hz and 90% spatial contrast. The stimuli were adjusted until the F2 (double frequency) response cycles were equalized, and conducted along both horizontal and vertical axes. All subsequent stimuli were aligned with the receptive field center. To determine the size of the receptive field center or surround of the RGC, a classical difference of the Gaussian model was applied to the area summation curve in response to uniform spots of varying diameters. The model is described as follows:

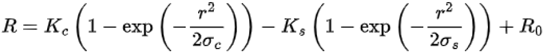

Whereas R is the response of the cell, *R*_0_ is the spontaneous activities, r is the radius of the expanding spot, *σ*_*C*_ and *σ*_*S*_ are the radius of the center, and surround, respectively, and *K*_*C*_ and *K*_*S*_ describe the scaling strength of the center and surround components respectively.

Naturalistic stimuli were generated using images from the van Hateren natural image database (van Hateren and van der Schaaf, 1998). For each image, we scaled the stimuli such that the brightest point in the entire image (from which the image patches were derived) was set as the highest monitor intensity. Natural image patches were generated from random samples of a given image. The background intensity when presenting natural image patches to cells was similarly defined as the mean intensity across the entire image from which the patches were drawn. Subsets of natural image patches were selected as stimuli, as described before (Turner and Rieke, 2016; Turner et al., 2018). In short, we incorporated the nonlinear subunit model previously developed for classifying image patches by their response nonlinearity. This involved comparing the difference in responses between spatially linear versus nonlinear RF models. We then selectively sampled across the full spectrum of image patches to present a balanced range, from highly nonlinear (with pronounced spatial contrast) to very linear (with minimal spatial contrast). This approach enabled us to efficiently investigate the entire spectrum of spatial contrasts found in natural images in a single experimental setup.

To construct the linear equivalent disc stimulus, we computed the weighted average pixel intensity of each image patch with a circular Gaussian function; Gaussian weights were determined for each recorded cell from responses to spots of a number of sizes. The weights for the Gaussian function were derived from each recorded cell’s response to differently sized spots. The Gaussian function’s width, defined by its two-standard deviation span, corresponded to the diameter of the aperture overlaying the image, which matched the measured size of the RF center. These natural image patches were then displayed at a retinal scale of 6.6 µm/pixel.

### Cell Identification and Selection Criteria

We identified alpha RGCs based on their characteristic large soma size and distinctive spike responses to light steps (Krieger et al., 2017). RGCs recordings were performed with retinal pigment epithelium (RPE) attached to preserve natural retinal function. To explore the underlying neural circuitry, we conducted recordings from multiple cell types: off-cone bipolar cells, rod bipolar cells, and A2 amacrine cells. For improved optical access, we targeted bipolar cells in flat-mount retina preparations with the RPE removed. We recorded from A2 amacrine cells with the RPE left intact. Cell types were distinguished based on their specific locations within the inner nuclear layer (INL) of the mouse retina and their unique electrophysiological signatures. In some cases, cell identities were further confirmed through dye fills. This comprehensive approach allowed us to examine the contributions of various cell types within the parallel pathways that shape αRGC responses to spatial structures.

### Center-Surround Subunit Model

The center-surround subunit model was implemented in MATLAB (adapted from (Turner et al., 2018)), to simulate the responses of retinal ganglion cells (RGCs) to grating stimuli of varying widths (Figure 4). The receptive field (RF) of each subunit was modeled as a difference of

Gaussians (DoG) with a variable inhibitory subunit center size *σ*_*inh*_ and surround *σ*_*inh*_ ^*surround*^ and the strength of the subunit surround (*δ*) was varied relative to the center (0.2 for weak surround, 0.5 for moderate surround, and 0.9 for strong surround, Figure 4). The equation for each subunit’s RF is:

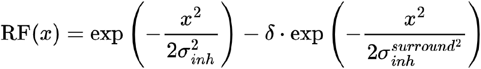

where x represents spatial position. Subunit responses were weighted by their position within the overall RF center (σ = 50 µm). Bar stimuli of varying widths were generated and circularly shifted across trials to simulate variability. This setup allowed us to examine how varying the subunit surround strength and size influenced spatial integration in the RGCs.

### Dynamic Synapse Model

We implemented a hybrid linear–nonlinear (LN) model that captures the core circuit dynamics we observed experimentally. The model incorporates two key elements: (1) an excitatory pathway governed by short-term synaptic depression, reflecting the bipolar cell terminals that undergo activity-dependent modulation, and (2) a static inhibitory pathway representing the A2 amacrine cell inputs. In this formulation, the excitatory output is regulated by a vesicle occupancy variable (representing available neurotransmitter vesicles at bipolar cell terminals) that changes according to synaptic depression dynamics.

The model accommodates both spatiotemporal (Figure 9) and hemifield-based stimuli (Figure 8). In the spatiotemporal configuration, each excitatory subunit integrates visual input across space and time, while receiving pooled inhibition from neighboring inhibitory subunits. For hemifield stimuli (e.g., pair-pulse and step paradigms, Figure 8),—spatial structure is collapsed, and each hemifield contains one excitatory and one inhibitory subunit. In this case, each excitatory subunit is modulated by the combined inhibition from both inhibitory subunits. (Model code is available at: https://github.com/chrischen2/spatialIntegration.git)

### Inhibitory Subunit: Static LN Model

Each inhibitory subunit *j* located at position *y*_*j*_ processes stimulus input *S*(*x,t*) through a spatial-temporal linear kernel *k*_*inh*_(*x*, τ), followed by a rectifying nonlinearity:

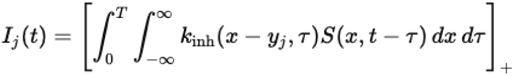

In the spatial-temporal model, the spatial receptive field of inhibitory subunits RF_inh_(x) is defined by the center-surround structure (with strong and local surround) characterized in Figure 4.

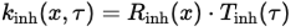

Where *T*_*inh*_ (τ) is the temporal filter of inhibitory subunits, which is extracted from experimental data. [·]_+_: half-wave rectification (threshold-linear nonlinearity). *R*_*inh*_ (*x*) is the spatial receptive field of inhibitory subunits, modelled as difference of gaussian (as Figure 4) and defined as equation below:

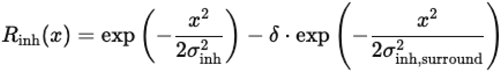

where σ_inh_ and σ_inh, surround_ denote the center and surround spatial scales of inhibitory subunit, respectively, and δ is the relative surround strength (dimensionless, 0<δ≤1). These parameters were chosen to match the spatial inhibitory receptive fields observed experimentally and illustrated in Figure 4.

In the hemifield-based temporal model (used in Figure 8), spatial integration is omitted and each hemifield is represented by a single inhibitory subunit. In that case:

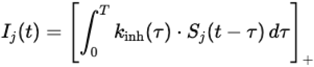

Here, *I*_*j*_ (*t*) is a static inhibitory signal and does not undergo dynamic synaptic changes. *S*_*j*_ (*t*) is the stimulus presented to hemifield *j*, where *j* ∈{left,right} indicates the hemifield.

### Excitatory Subunit: Dynamic LN Model with Presynaptic Inhibition

Excitatory drive for each excitatory subunit *i*, located at *x*_*i*_, receives linearly filtered input via an excitatory kernel *k*_*exc*_ (*x*, τ), followed by presynaptic inhibition with strength factor α:

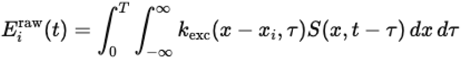

This signal is modulated by presynaptic inhibition, pooled from nearby inhibitory subunits using a Gaussian weighting function:

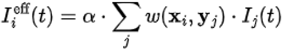

Where *w*(*x*_*i*_, *y*_*j*_) defines the spatial influence of inhibitory subunit j on excitatory subunit i, and α is a global inhibition strength factor.

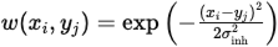

The presynaptically modulated excitatory input is passed through a static piecewise linear nonlinearity Φ(·) defined as:

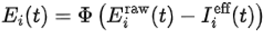

In our model, excitation output nonlinearity Φ(x) is given by:

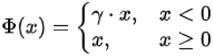

where γ, or the rectification ratio, ∈(0,1) is a constant that scales the slope of the negative portion of the input.

Then, the positive portion of this signal is dynamically scaled by the vesicle occupancy ni(t)∈[0,1], which captures short-term synaptic depression. The final output of subunit i is:

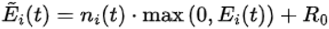

This formulation scales output by the availability of synaptic vesicles. The dynamics of vesicle occupancy are given by:

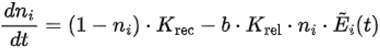

where *K*_*rec*_ and *K*_*rel*_ are vesicle recovery and release rates, and b is a gain constant.

In the temporal (hemifield) model, each hemifield contains one excitatory and one inhibitory subunit, and inhibitory pooling reduces to:

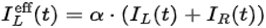

### Pooling onto the Ganglion Cell

In the spatiotemporal model, the total excitatory and inhibitory drive to the ganglion cell is obtained by spatially weighted summation across subunits. Let *W*_*exc*_ (*x*_*i*_) *and W*_*inh*_ (*y*_*j*_) be

Gaussian spatial weights centered on the soma:

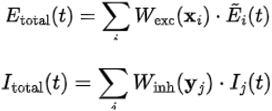

For hemifield-based stimuli (e.g., step and pair-pulse paradigms), each hemifield consists of one excitatory and one inhibitory subunit. Spatial weighting is omitted, and the total excitatory and inhibitory drives computed as the average of the subunit outputs.

### Model Parameters

For all simulations, constant parameters—such as the vesicle recovery and release rates (*K*_rec_,*K*_rel_, release gain (*b*). The rectification ratio γ, which defines the slope of the subthreshold (negative) region of the excitatory nonlinearity Φ(x), was constrained using experimental data. Specifically, we used values derived from nonlinearities measured in Off-transient alpha RGCs stimulated with temporal Gaussian noise and sinusoidal modulations under mesopic conditions.

To further constrain the model,the initial vesicle occupancy *n*_0_ was not treated as a free parameter but was **i**nferred directly from experimental measurements of baseline excitation.

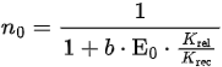

This expression arises from setting 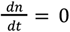 in the vesicle dynamic equation under constant baseline drive and we also assume in the initial steady state baseline inhibition is zero. By using this constraint, the model ties the initial condition *n*_0_ directly to measurable physiological quantities.

The qualitative dynamics we study arise robustly from the model structure rather than specific parameter tuning.

**Supplementary Table 1.** Parameters for the Temporal and Spatial-Temporal Dynamic Synapse Models.

| Parameter | Description | Temporal Only Model Value | Spatial-Temporal Model Value | Unit |
| --- | --- | --- | --- | --- |
| $K_{rec}$ | Vesicle recovery rate | 10 | 10 | Hz |
| $K_{rel}$ | Vesicle release rate | 5 | 5 | Hz |
| $b$ | Release gain | 5 | 5 | - |
| $\gamma$ | Rectification ratio (exc) | 0.3 | 0.3 | - |
| $E_0$ | Norm. excitation baseline | 0.3 | 0.3 | - |
| $\alpha$ | Presynaptic inhibition Strength [0,1] | 0.9 | 0.9 | - |
| $\beta$ | Sensitivity of release rate to inhibition | 1 | 1 | - |
| $E_0$ | Baseline excitation | 0.3 | 0.3 | - |
| $\sigma_{exc}$ | Excitatory subunit center size | - | 22 | $\mu m$ |
| $d_{exc}$ | Excitatory subunit spacing | - | 44 | $\mu\text{m}$ |
| $\sigma_{pool,exc}$ | Excitatory pooling radius | - | 50 | $\mu\text{m}$ |
| $\sigma_{inh}$ | Inhibitory subunit center size | - | 12 | $\mu\text{m}$ |
| $d_{inh}$ | Inhibitory subunit spacing | - | 24 | $\mu\text{m}$ |
| $\sigma_{inh,surround}$ | Inhibitory surround size | - | 20 | $\mu\text{m}$ |
| $\delta$ | Surround strength onto inhibition subunit | - | 0.9 | - |
| $\sigma_{pool,inh}$ | Inhibitory pooling radius | - | 100 | $\mu\text{m}$ |

## Acknowledgements

We thank S. Cunnington for technical support and Wei Wei and Mike Manookin for their insightful feedback to the manuscript. This work was supported by NIH EY028541.

